# MOF-mediated Histone H4 Lysine 16 Acetylation Governs Mitochondrial and Ciliary Functions By Controlling Gene Promoters

**DOI:** 10.1101/2022.11.23.517702

**Authors:** Dongmei Wang, Haimin Li, Navdeep S Chandel, Yali Dou, Rui Yi

## Abstract

Histone H4 lysine 16 acetylation (H4K16ac), governed by the histone acetyltransferase (HAT) MOF, orchestrates critical functions in gene expression regulation and chromatin interaction. However, how does MOF and H4K16ac control cellular function and regulate mammalian tissue development remains unclear. Furthermore, whether the function of MOF is mediated by MSL or NSL, two distinct MOF-containing HAT complexes, have not been determined during mammalian development. Here we show that conditional deletion of *Mof* but not *Kansl1*, an essential component of the NSL complex, causes severe defects during murine skin development. In the absence of *Mof* and H4K16ac, basal epithelial progenitors of mammalian skin fail to establish the basement membrane and cell polarity, causing the failure of self-renewal. Furthermore, epidermal differentiation and hair growth are severely compromised, leading to barrier defects and perinatal lethality. Single-cell and bulk RNA-seq, in combination with MOF ChIP-seq, reveal that MOF regulated genes are highly enriched in mitochondria and cilia. Mechanistically, MOF coordinates with RFX2 transcription factor, which preferentially binds to gene promoters, to regulate ciliary and mitochondrial genes. Importantly, genetic deletion of *Uqcrq*, a nuclear-encoded, essential subunit for electron transport chain (ETC) Complex III, recapitulates the defects of epidermal differentiation and hair follicle growth observed in MOF cKO. Together, this study reveals the requirement of MOF-mediated epigenetic mechanism for mitochondria and cilia, and demonstrates the important function of the MOF/ETC axis for mammalian skin development.

## Main

Histone H4 lysine 16 acetylation (H4K16ac) has been implicated to play important functions, ranging from chromatin decompaction, gene expression regulation to priming future gene activation^1–6^. Although histone acetylation is generally associated with gene expression activation, H4K16ac is distinct from the acetylation at positions 5, 8 and 12 of histone H4 with specific transcriptional outcomes when ablated in budding yeast^7^. Structural and biophysical studies demonstrate that H4K16ac plays an essential role in transcription activation mediated by its ability to regulate both nucleosome structure and interaction with chromatin-binding proteins, such as epigenetic modifiers for other histone marks^8–10^.

H4K16ac is catalyzed by the MYST-family lysine acetyltransferase (KAT) MOF^11,12^ (also known as KAT8), which is broadly conserved in fly, mouse and human. However, MOF has been detected in two highly conserved and mutually exclusive complexes, MSL (male-specific lethal) and NSL (non-specific lethal), in fly, mouse and human cells^11,13–15^. Whereas MSL is largely specific to H4K16ac^16,17^, recent studies in mammalian cells have demonstrated that many functions of MOF, including promoting mtDNA transcription^1^, are mediated by NSL, which has a broader range of substrates than H4K16ac^13,16,18^. While these studies have demonstrated complex functions of MOF in mammalian cells^3,19–21^, the lack of genetic study for MOF comparatively with the NSL complex has severely hindered the understanding of MOF/H4K16ac-mediated functions in mammalian development.

Furthermore, it remains unclear how MOF-regulated gene expression program governs cellular functions and lineage differentiation during mammalian tissue development. Mechanistically, although H4K16ac has the unique ability to decompact chromatin structure and orchestrate other chromatin marks to activate gene expression^7,9,10,22^, whether specific transcription factors (TFs) require MOF and H4K16ac to regulate gene expression and cellular functions remains unknown.

To address these important questions, we leverage spatiotemporally well-established lineage specification and tissue morphogenesis in mammalian skin to define the function and mechanism of MOF and H4K16ac by using epithelial specific conditional knockout (cKO) mouse models for both *Mof* and *Kansl1*, an essential component of NSL^14,15,23^. We provide genetic evidence that *Mof* but not *Kansl1* plays an important role in embryonic skin development. We have further discovered that MOF-regulated genes are highly enriched in genes involved in two important organelles, mitochondria and primary cilia. Notably, many MOF targets, in particular ciliary genes and, to a lesser extent, mitochondrial genes, are regulated by a family of conserved RFX (Regulatory Factor binding to the X-box) TFs preferentially through the promoters in a MOF-dependent manner. Finally, we provide genetic evidence that MOF-regulated mitochondrial functions, through the electron transport chain (ETC), are critical for epidermal differentiation and hair follicle development.

## Results

### MOF and H4K16ac are required for embryonic skin development

We first examined the functions of MOF during embryonic skin development by conditionally deleting MOF (cKO) with an epithelial specific Cre line (*Krt14-Cre*)^5,24^.

Immunofluorescence (IF) staining for H4K16ac, which requires the acetyltransferase activity of MOF^23^, in embryonic skin revealed that this histone mark is ubiquitously present in both epithelial and dermal cells of the skin. Homozygous deletion of *Mof* by *Krt14-Cre* (MOF cKO) resulted in a gradual depletion of H4K16ac in embryonic epidermis (Fig. 1a). At embryonic day 15.5 (E15.5), H4K16ac level was reduced to ∼30% of control. By E16.5, H4K16ac level was further reduced to <20%. In contrast, H4K16ac signal in dermis was intact.

**Fig. 1.**
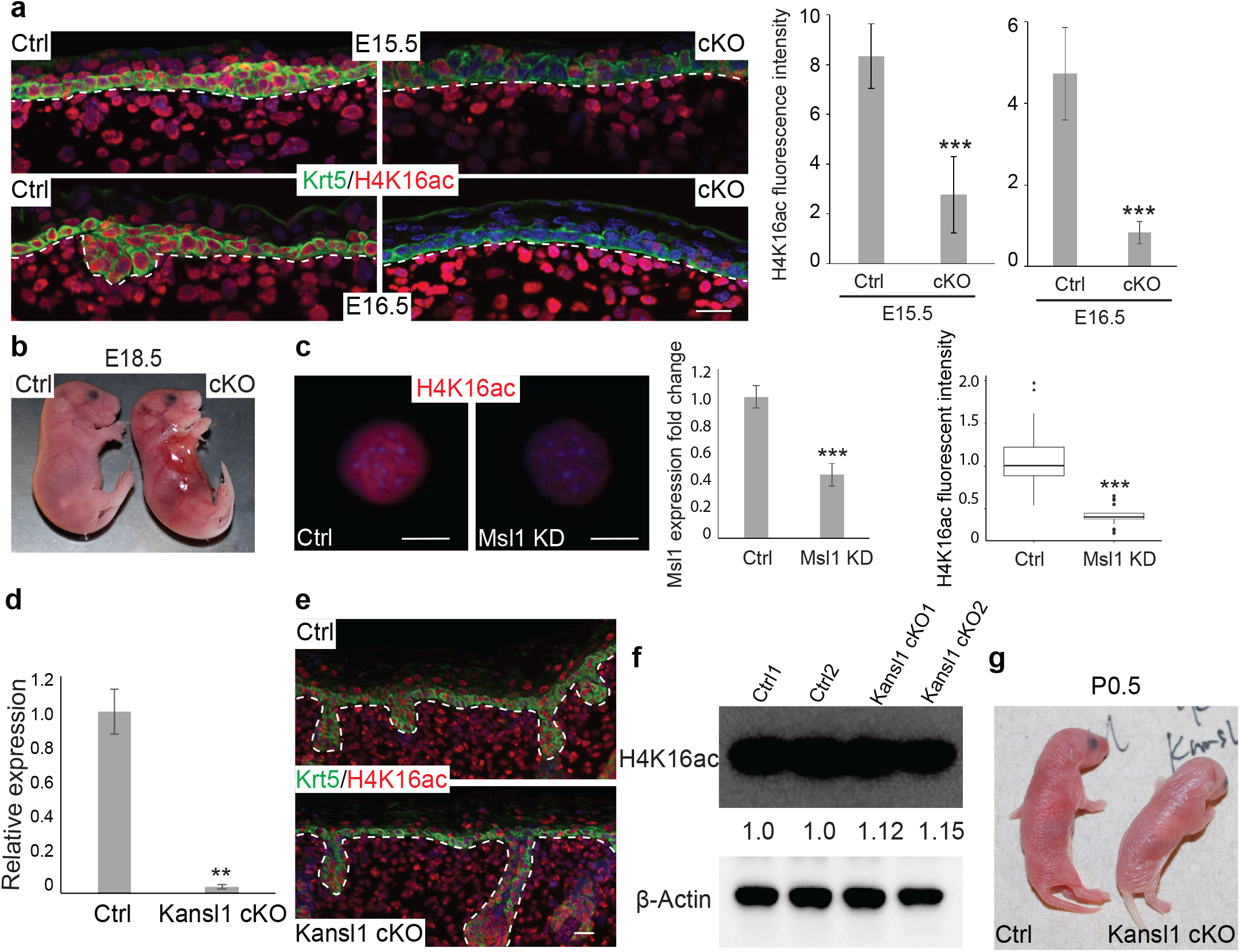
MOF and H4K16ac are required for embryonic skin development. **a**, Immunofluorescence staining showing ubiquitous distribution of H4K16ac throughout the embryonic skin in control (Ctrl) animals, and gradual loss of H4K16ac in the epidermis of MOF/K14Cre conditional KO (cKO) animals. **b**, Partial loss of skin and thinner skin in cKO animals at E18.5. **c**, IF staining and quantification showing reduced H4K16ac upon knockdown of Msl1 by shRNA in mouse keratinocytes. **d**, Knockout of Kansl1 exon3 in epidermis confirmed by qPCR. **e**, H4K16ac staining in Kansl1 cKO. **f**, Quantification of H4K16ac level by Western Blot in Kansl1 cKO epidermal cells. **g**, Indistinguishable appearance of Kansl1 cKO animals with control. Ctrl, control. White dashed lines mark epidermal-dermal boundary (**a** and **e**). *P* values were calculated by unpaired two-tailed Student’s *t* test (**a, c** and **d**), **, *P* < 0.01; ***, *P* < 0.001. Scale bar, 20 μm (**a** and **e**), 10 μm (**c**).

Although MOF cKO mice were born at the expected Mendelian ratio and had a similar size to their wildtype (WT) or heterozygous (het) littermate controls, they died within a few hours after birth, indicative of severe defects in the skin. Indeed, most newborn cKOs missed parts of their skin, usually at the flank near the forelimbs. To circumvent skin detachment caused by mechanical stress during birth, we collected embryos at E18.5. Yet, MOF cKO embryos already developed fragile skin that was occasionally detached from the underlying dermis (Fig. 1b). In areas where the skin was present, it was very thin, indicating defective epidermal formation. These data provided initial evidence that loss of MOF compromised epidermal adhesion and differentiation during embryonic skin development.

MOF resides in mutually exclusive and evolutionarily conserved MSL and NSL complexes. Although both complexes are capable of acetylating H4K16, MSL has been identified as the main HAT acetylating H4K16^16,20,25^. In contrast, non-histone acetylation activities, such as acetylation of p53 and LSD1^13,16,18^ as well as intramitochondrial gene regulation^1^, are restricted to NSL. Previous studies have further indicated that NSL is responsible for the broader acetylation activities of MOF on H4K5 and H4K8^16,20^. To determine whether MOF functions through NSL or MSL in skin epithelial cells, we first used shRNA to knockdown *Msl1*, an essential component of MSL^16,23^, in primary mouse keratinocytes. As expected, downregulation of *Msl1* resulted in significantly reduced H4K16ac (Fig. 1c).

To distinguish whether the severe defects observed in MOF cKO are mediated by deacetylation of H4K16 or other substrates of MOF mediated by NSL, we deleted an essential component of NSL, *Kansl1* also known as MSL1v1^13,15^, in epithelial cells of the skin by using the same *Krt14-Cre* (Kansl1 cKO). The knockout strategy removes the third exon of *Kansl1* (Extended Data fig. 1a), which generates two consecutive premature stop codons immediately in the next exon and abrogates the expression of well-characterized functional domains of KANSL1, including the WDR5 binding domain and MOF binding domain^15^.

Epithelial cell specific deletion was confirmed by genotyping (Extended Data fig. 1b), qPCR (Fig. 1d) and RNA-seq (Extended Data fig. 1c). Genetic deletion of *Kansl1* in the epidermis, however, did not strongly affect H4K16ac as shown by IF staining (Fig. 1e) and western blot (Fig. 1f). In sharp contrast to the severe defects observed in MOF cKO mice, *Kansl1* cKO mice were indistinguishable from control littermates (Fig. 1g) and lived normally. Upon closer examination, a slight increase of apoptosis was detected in *Kansl1* cKO epidermis (Extended Data fig. 1d). Basement membrane (BM) marked by β4 integrin, epidermal differentiation marked by KRT1, hair follicle progenitors marked by SOX9 and cell proliferation marked by Ki67 all appeared normal and indistinguishable between control and *Kansl1* cKO (Extended Data fig. 1e). Furthermore, RNA-seq of control and *Kansl1* cKO epidermal samples only identified 51 downregulated genes and 35 upregulated genes in *Kansl1* cKO (Extended Data fig. 1f), consistent with the minimum perturbation to skin development. Taken together, these data establish the requirement of MOF-controlled H4K16ac during embryonic skin development and reveal the dispensable role of the NSL complex for MOF-mediated functions in the skin.

### Loss of MOF compromises basal cell adhesion, epidermal differentiation and hair follicle morphogenesis

We next performed in-depth examination of epidermal defects in MOF cKO skin, which tracked closely with H4K16ac depletion. At E16.5 when H4K16ac was largely depleted in the cKO epidermis, epithelial layers of MOF cKO skin started to show signs of reduced adhesion and had fewer differentiated layers (Fig. 2a). By E18.5, epidermal detachment became widespread, and the reduction of basal cells and compromised epidermal differentiation, shown by reduced granular layers and stratum corneum, were evident in morphological analysis (Fig. 2a). To capture the early defects concurrently with the loss of H4K16ac, we focused our analyses on E16.5 samples. To determine molecular features associated with epidermal detachment, we examined basal epithelial cells for the integrity of the BM. In MOF cKO, β4 integrin and collagen XVII, two key components of the BM, were detected in KRT5+ basal cells but failed to localize to the BM (Fig. 2b). Electronic microscopy (EM) revealed a strong reduction of hemidesmosome and, in some regions, a complete loss of the BM (Fig. 2c), corroborating the disorganized patterns of β4 and collagen XVII. Properly formed BM not only promotes epithelial adhesion but also allows the establishment of basal cell polarity, which, in turn, permits epidermal stratification and differentiation^26^. Consistent with the severely compromised BM, MOF cKO basal cells also failed to establish polarity, revealed by disoriented Pericentrin signals (Fig. 2d) and disorganized adherens junctions (Fig. 2e).

**Fig. 2.**
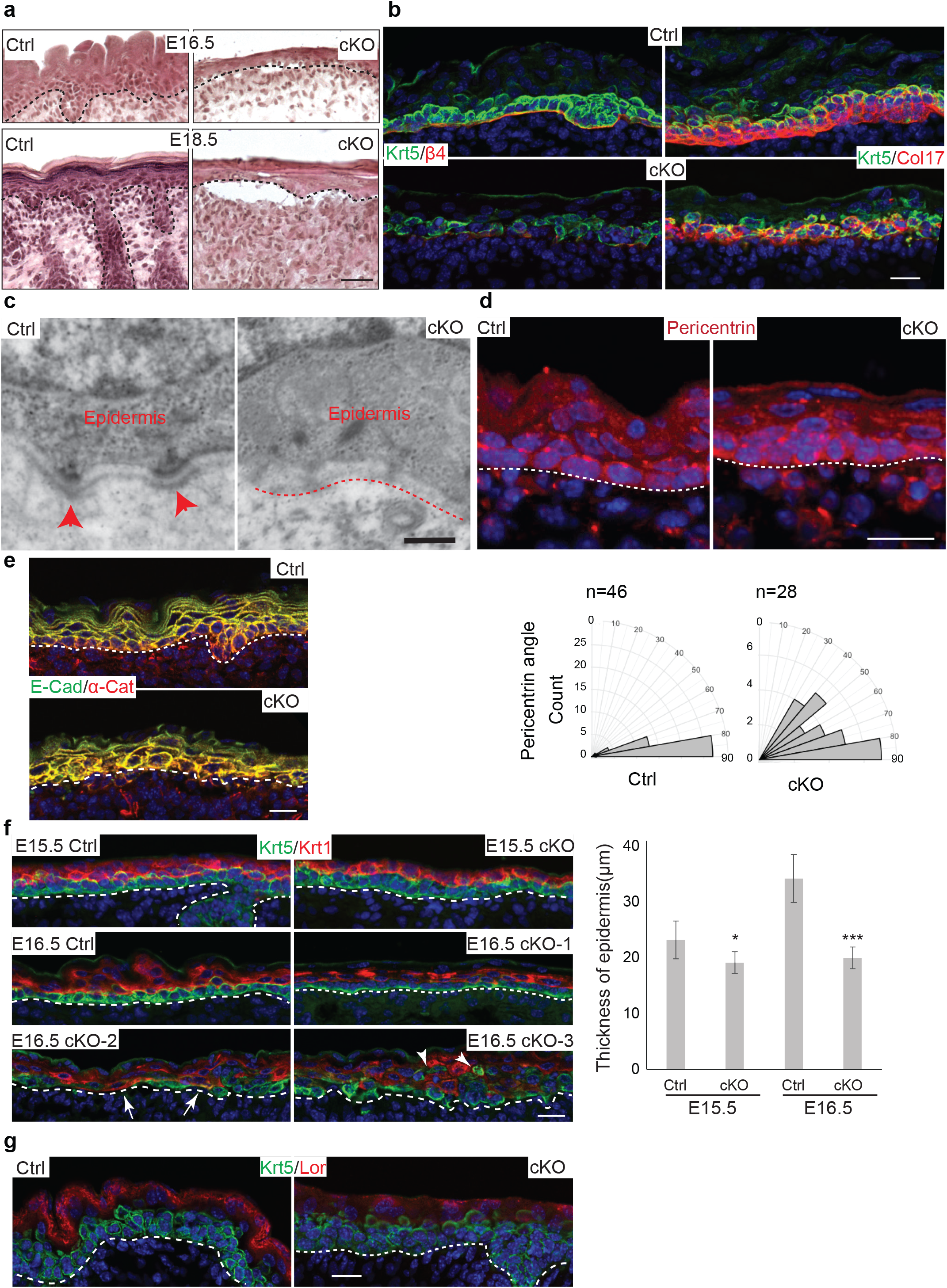
Loss of MOF compromises basal cell adhesion, epidermal differentiation and hair follicle morphogenesis. **a**, HE staining at E16.5 and E18.5 showing detachment of epidermis from dermis in cKO. **b**, Compromised basement membrane in cKO as shown by β4 integrin and collagen XVII (Col17) staining. **c**, Loss of hemi-desmosome and basement membrane shown by electronic microscopy. Red arrowheads in Ctrl indicate hemi-desmosome, red dashed line in cKO indicates missing basement membrane. **d**, Disorientation of basal cells indicated by pericentrin staining. Quantification is their angles relative to the basement membrane. n indicates number of cells counted. **e**, Loss of polarity in basal cells as shown by distribution of adheres junction markers E-Cadherin (E-Cad) and α-Catenin (α-Cat). **f**, Thinner epidermis in cKO shown by co-staining of basal layer marker Krt5 and the spinous layer marker Krt1. Arrows indicate Krt1^+^ cells came in contact with basement membrane, arrowheads point to delaminated Krt5^+^ cells that did not acquire Krt1 expression. **g**, Defect in terminal differentiation indicated by granular layer marker Loricin (Lor) staining. Ctrl, control. Black (**a**) and white (**d, e, f** and **g**) dashed lines mark epidermal-dermal boundary. *P* values were calculated by unpaired two-tailed Student’s *t* test (**f**), *, *P* < 0.05; ***, *P* < 0.001. Scale bar, 50 μm (**a**), 20 μm (**b, d, e, f** and **g**), 200 nm (**c**).

Instead of apical and lateral localization of E-Cadherin in control basal cells, E-Cadherin mis-localized to the basal side, reflecting lost polarity and compromised adhesion to the BM. The strongly compromised epithelial cell adhesion and BM prompted us to examine whether epithelial fate specification was altered in the absence of MOF and H4K16ac. However, the robust expression of KRT5 and ΔNp63 (Extended Data fig. 2a), the master TF required for epidermal fate specification^27^, ruled out the possibility of failed epidermal fate in MOF cKO.

We next examined epidermal differentiation. At E15.5 prior to the significant depletion of H4K16ac (Fig. 1a), both KRT5+ basal and KRT1+ spinous layers were largely intact in MOF cKO epidermis although the overall thickness of the epidermis was slightly reduced in cKO (Fig. 2f). However, by E16.5, whereas control epidermis continued to thicken with the expansion of differentiated spinous and granular layers, MOF cKO epidermis had much thinner spinous and granular layers (Fig. 2f-g). In addition, basal cell proliferation was reduced, and the detached epidermal regions showed the strongest reduction (Extended Data fig. 2b). Apoptotic cells, marked by activated Caspase 3, were also detected mostly in the basal layer of the cKO epidermis (Extended Data fig. 2c). Furthermore, reduced cell adhesion to the BM caused premature delamination of basal cells. Many of these delaminated cells were stalled to a KRT5+/KRT1-basal-like phenotype (Fig. 2f, arrowheads), failing to differentiate. Collectively, defects in cell adhesion and differentiation severely compromised epidermal integrity in the suprabasal layers (Fig. 2f). Consistent with defective epidermal differentiation, granular layer, marked by Loricrin (Lor), was also thinner and had weaker Lor signals in MOF cKO (Fig. 2g). Finally, KRT6 was robustly detected in the suprabasal layers of MOF cKO epidermis (Extended Data fig. 2d), reflecting a stressed cellular state.

We then turned to hair follicle (HF) induction and morphogenesis. Overall, the number of morphologically discernible HFs were significantly reduced (Fig. 2a). In the few HFs that were still present in cKO, however, the stem cell/progenitor marker, SOX9, was readily detected (Extended Data fig. 2e). This argued against the complete failure of HF fate specification. Indeed, LEF1, the *bona fide* TF for the activated WNT signaling pathway and the marker for hair placode formation^28,29^, was readily detected in regularly spaced hair placodes in MOF cKO skin (Extended Data fig. 2f). These data suggest that HF fate was induced but the downward growth of HFs was arrested in the absence of MOF and H4K16ac, reminiscent of the hair growth defects observed in Shh KO^30,31^. Altogether, these data illustrate widespread impact of MOF and H4K16ac on epidermal and HF cell lineages.

## Single-cell RNA-seq reveals widespread downregulation of nuclear encoded mitochondrial genes

We next performed single-cell RNA-seq (scRNAseq) to interrogate the changes of transcriptome and cellular states caused by the deletion of *Mof* at E16.5 when H4K16ac was depleted. Dorsal skin samples from two pairs of control and MOF cKO embryos were profiled with the 10x Chromium Single Cell 3’ kit. After quality control, 5,656 cells were recovered, and ten major cell types were identified in control and cKO samples, including epithelial cells, dermal cells and immune cells among others (Fig. 3a). In two control samples, epithelial cells represented 17% and 20% of the total population, respectively. In contrast, in two cKO samples, epithelial cells represented only 5% and 2.7% of the total, respectively. These results were consistent with the reduced epithelial cell populations as observed in our phenotypical analysis. In addition, dermal cell clusters from cKO samples were slightly deviated from control although MOF was only deleted from epithelial cells and H4K16ac was intact in the dermis (Fig. 3a and Extended Data fig. 3a), likely reflecting the response of dermal cells to MOF cKO epithelium. Gene ontology (GO) enrichment analysis for upregulated genes in MOF cKO dermis revealed that altered collagen fibril organization and increase inflammatory response contributed to the different cellular states (Fig. 3b). MOF cKO samples also had more immune cell infiltration (8.4% in cKO vs 3.7% in Ctrl), consistent with the increased inflammatory signatures in the dermal cells.

**Fig. 3.**
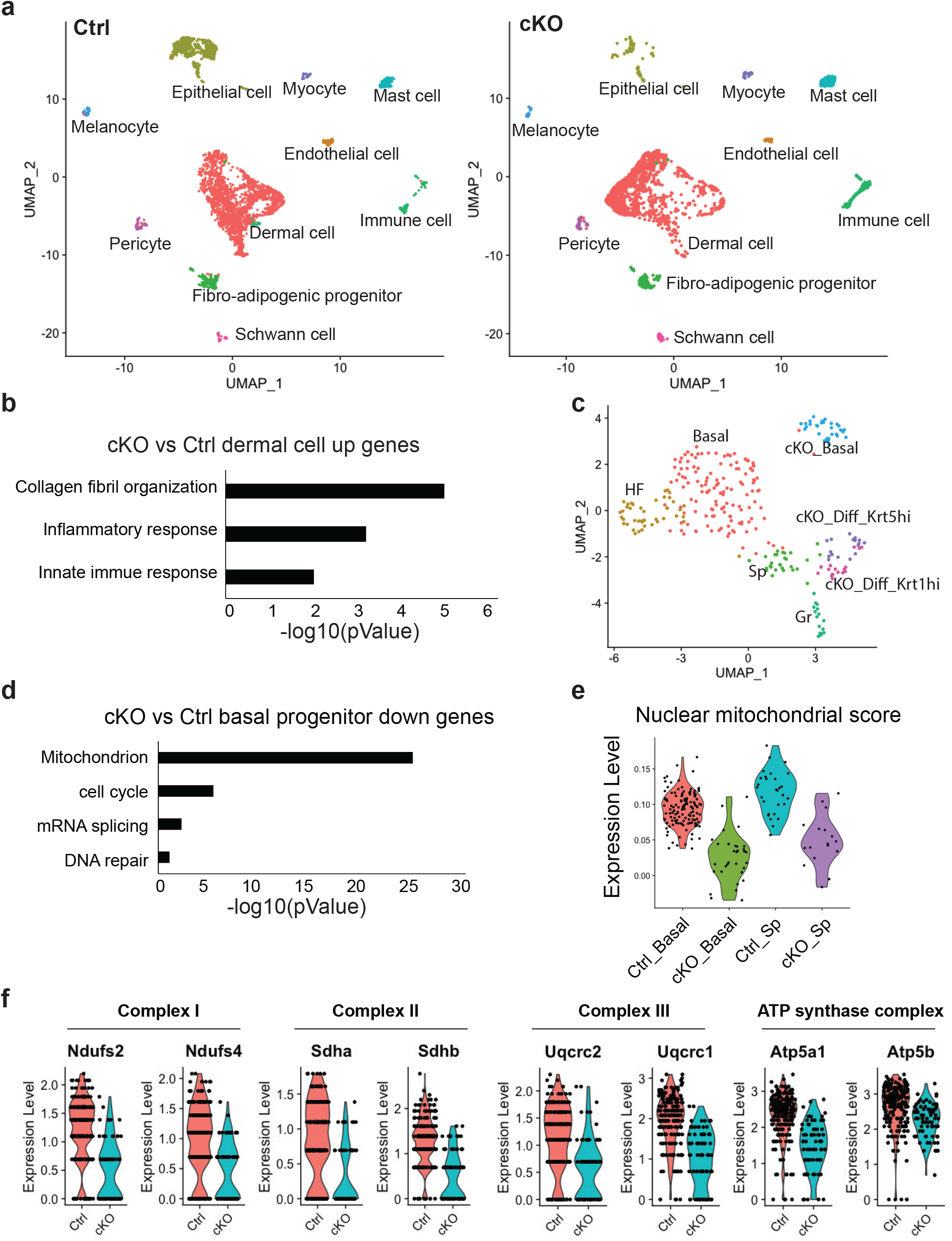
Single-cell RNA-seq reveals widespread downregulation of nuclear encoded mitochondrial genes. **a**, Ten distinct cell types detected in E16.5 control and cKO dorsal skin samples by scRNAseq. **b**, Gene ontology (GO) terms of up-regulated genes in cKO dermal cells. **c**, UMAP clustering of epidermal cells. Four populations were identified in control and three in cKO. Basal, basal progenitor cells; HF, hair follicle cells; Sp, spinous layer cells; Gr, granular layer cells; cKO_Basal, basal like progenitor cells in MOF cKO; cKO_Diff_Krt5hi, Krt5^hi^Krt1^low^ suprabasal cells in MOF cKO; cKO_Diff_Krt1hi, Krt5^low^Krt1^hi^ suprabasal cells in MOF cKO. **d**, GO terms of down-regulated genes in MOF cKO *vs*. control basal progenitor comparison. **e**, Aggregated nuclear-encoded mitochondrial gene expression score calculated using 1,158 mitochondrial genes. **f**, Reduced expression of electron transport chain complex genes in cKO. Ctrl, control.

Next, we focused on four epithelial clusters that were detected in control, including basal epidermal progenitor (marked by high Krt14 and Krt5 and high proliferation), spinous layer of the epidermis (marked by Krt1 and Krt10), granular layer of the epidermis (marked by Lor and Ivl) and HF progenitors (marked by Sox9 and Pthlh) (Fig. 3c and Extended Data fig. 3b-c). Interestingly, only two major clusters, both of which were distinct from control clusters, were detected in MOF cKO (Fig. 3c and Extended Data fig. 3b), underscoring the profound changes in cKO transcriptome. One cKO cluster, which had high levels of Krt14 and Krt5 as well as other basal markers (Supplemental Table 1), represented the basal cell population of cKO (Fig. 3c and Extended Data fig. 3c). The other cKO cluster, representing differentiated epithelial cells, could be further separated into two sub-clusters based on differential Krt5 and Krt1 expression patterns (Fig. 3c and Extended Data fig. 3c). One Krt1^high^/Krt5^low^ sub-cluster probably corresponded to the differentiated spinous layer cells. The other Krt1^low^/Krt5^high^ sub-cluster was reminiscent of the KRT5+/KRT1-suprabasal cells seen in morphological studies (Fig. 2f). Thus, scRNAseq captured major cell populations and identified defective epidermal differentiation in control and cKO skin.

We next performed differential gene expression analysis of basal and suprabasal epithelial populations. In addition to reduced gene expression in BM and ECM (Extended Data fig. 3d), which was consistent with morphological analysis, downregulated genes in MOF cKO basal cells were highly enriched in GO categories of mitochondrion, cell cycle, mRNA splicing and DNA repair (Fig. 3d). The same GO categories for downregulated genes were also enriched in MOF cKO vs control suprabasal cells (Extended Data fig. 3e). In addition, genes associated with cornified envelope were the most significantly downregulated in MOF cKO suprabasal cells, confirming the strongly compromised epidermal differentiation (Extended Data fig. 3e).

We next examined the expression of nuclear-encoded mitochondrial genes in basal and suprabasal cells in control and MOF cKO. By using a mitochondrial gene expression score, which was calculated by using a curated list of 1,158 mitochondrial genes, we observed that their expression was elevated in differentiated suprabasal cells, compared with basal progenitors, in control. Furthermore, in both basal and suprabasal cells, the mitochondrial gene expression score was strongly downregulated in MOF cKO (Fig. 3e).

Strikingly, downregulated mitochondrial genes were highly enriched in the essential components of the electron transport chain (ETC) Complexes I, II, III and ATP synthase complex as well as mitochondrial matrix (Fig. 3f and Extended Data fig. 3f). Collectively, these data reveal strong changes of transcriptome and cellular states, caused by loss of MOF, during epidermal development.

### MOF cKO causes widespread downregulation of ciliary genes

MOF and H4K16ac have been implicated in the regulation of “housekeeping” genes, which are generally defined by their relatively high expression and essential functions across different cell types. In this regard, genes associated with mitochondrial functions, cell cycle, mRNA splicing and DNA repair, which were detected as top downregulated gene groups in our scRNAseq analysis, appeared to corroborate this notion. However, scRNAseq is limited to detect highly expressed transcripts. To comprehensively identify all genes that are regulated by MOF, we performed bulk RNAseq by using FACS purified epithelial cells isolated from control and MOF cKO skin samples at E16.5. Using FDR≤0.01 and 1.5x fold change as cut-off, we identified 1,734 downregulated genes and 1,752 upregulated genes (Fig. 4a). Consistent with scRNAseq data, important BM and ECM genes, such as *Col7a1, Frem2* and *Lamc1*, were significantly downregulated (Extended Data fig. 4a). Furthermore, nuclear-encoded mitochondrial genes, cell cycle and DNA repair genes were also significantly downregulated, mirroring the findings from scRNAseq. Surprisingly, genes associated with primary cilia emerged as the most highly enriched GO category among downregulated genes (Fig. 4b). Interestingly, most of these ciliary genes were not detectable in either control or cKO scRNAseq datasets because of their relatively low expression. To further examine the widespread downregulation of ciliary genes, we curated a list of 659 ciliary genes from the GSEA^32^, among which 443 genes were detected in the bulk RNAseq, and performed GSEA analysis. The result confirmed the global downregulation of ciliary genes in epithelial cells of MOF cKO (Fig. 4c). Upon closer inspection, these downregulated genes were broadly involved in both biogenesis and function of primary cilia^33^, including ciliary basal body, transition zone, intraflagellar transport and Shh signaling (Fig. 4d).

**Fig. 4.**
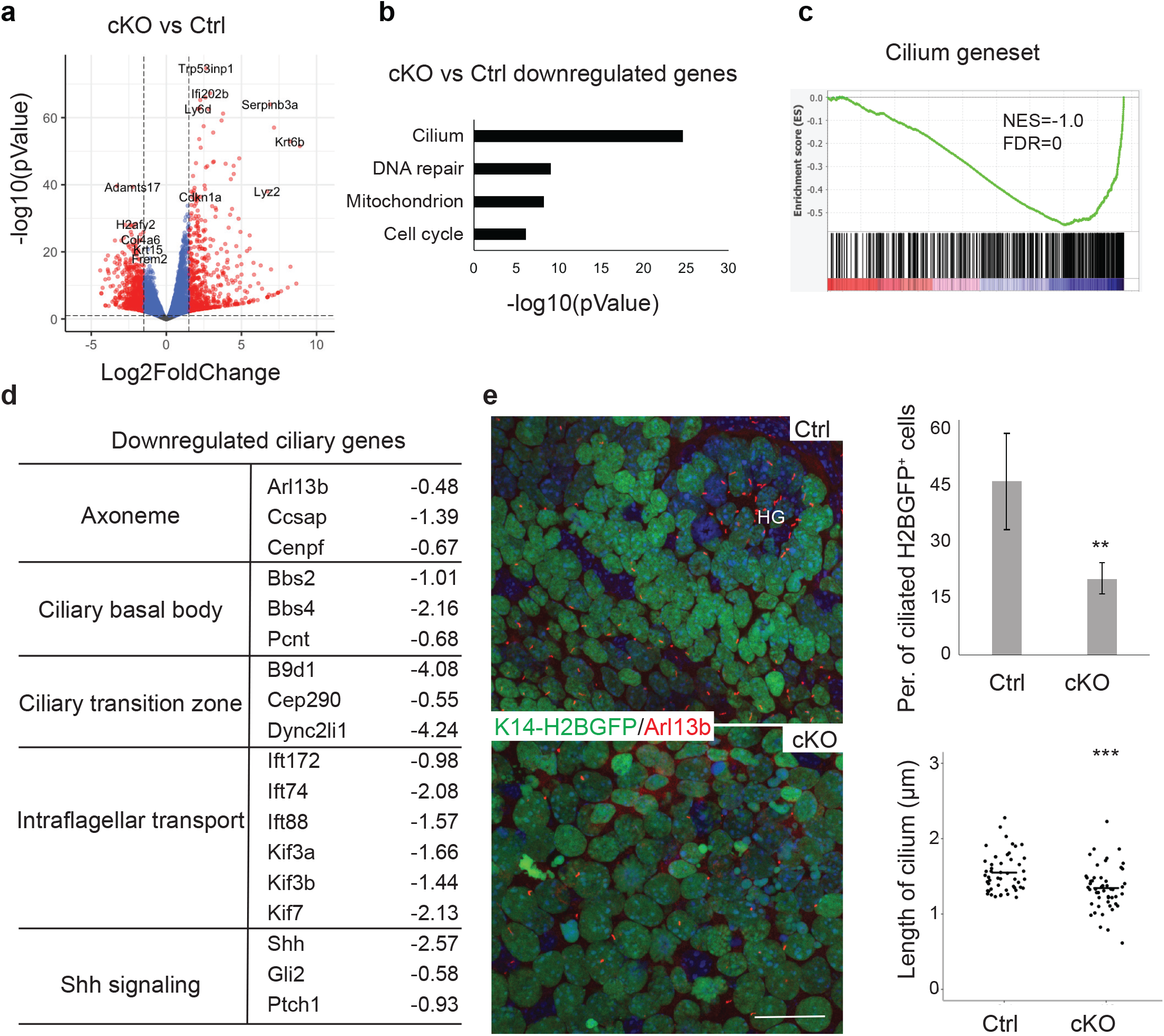
MOF cKO causes widespread downregulation of ciliary genes. **a**, Volcano plot of differentially expressed genes in cKO compared with control. Red dots represent genes showing fold change > 1.5 and FDR < 0.01. **b**, GO terms of down-regulated genes. **c**, GSEA for a set of 443 ciliary genes. **d**, Examples of down-regulated ciliary genes in different structures of a cilium. **e**, Reduced number of ciliated epithelial cells as well as reduced cilia length in cKO as marked by Arl13b. *P* value was calculated by unpaired two-tailed Student’s *t* test (**e**), **, *P* < 0.01; ***, *P* < 0.001. Scale bar, 20 μm (**e**).

These molecular findings prompted us to examine primary cilia in MOF cKO skin. In control, many primary cilia were detected in basal progenitors and in hair germs with the polarized localization, consistent with previous reports^34^. In contrast, primary cilia were strongly reduced in their numbers in MOF cKO. For the few cilia that persisted in MOF cKO, their length was significantly reduced (Fig. 4e). Together, these data reveal that numerous ciliary genes were downregulated in MOF cKO and uncover the requirement of MOF-regulated transcriptome for primary cilia in the skin.

### MOF regulates gene expression preferentially through promoters

We next investigated the mechanism of MOF-mediated regulation. We performed MOF ChIPseq and Cut&Run assays for H4K16ac, H3K4me3, H3K4me1 and H3K27ac by using freshly isolated epidermal keratinocytes from newborn animals. Consistent with previous studies, most peaks (3,100 peaks from the c1/c2/c3 clusters) co-occupied by MOF and H4K16ac were generally associated with active promoters and enhancers (Fig. 5a), illustrated by the locus of *Mof* and *Ndufs2* (Fig. 5b). We also detected 1,799 peaks from the c4 cluster, which were occupied by MOF, H4K16ac, H3K4me1 and H3K27ac but not H3K4me3. In addition, we also detected 5,915 peaks from the c5/c6 clusters that were occupied only by MOF but not H4K16ac or other marks, probably due to technical issues related to MOF ChIP and Cut&Run.

**Fig. 5.**
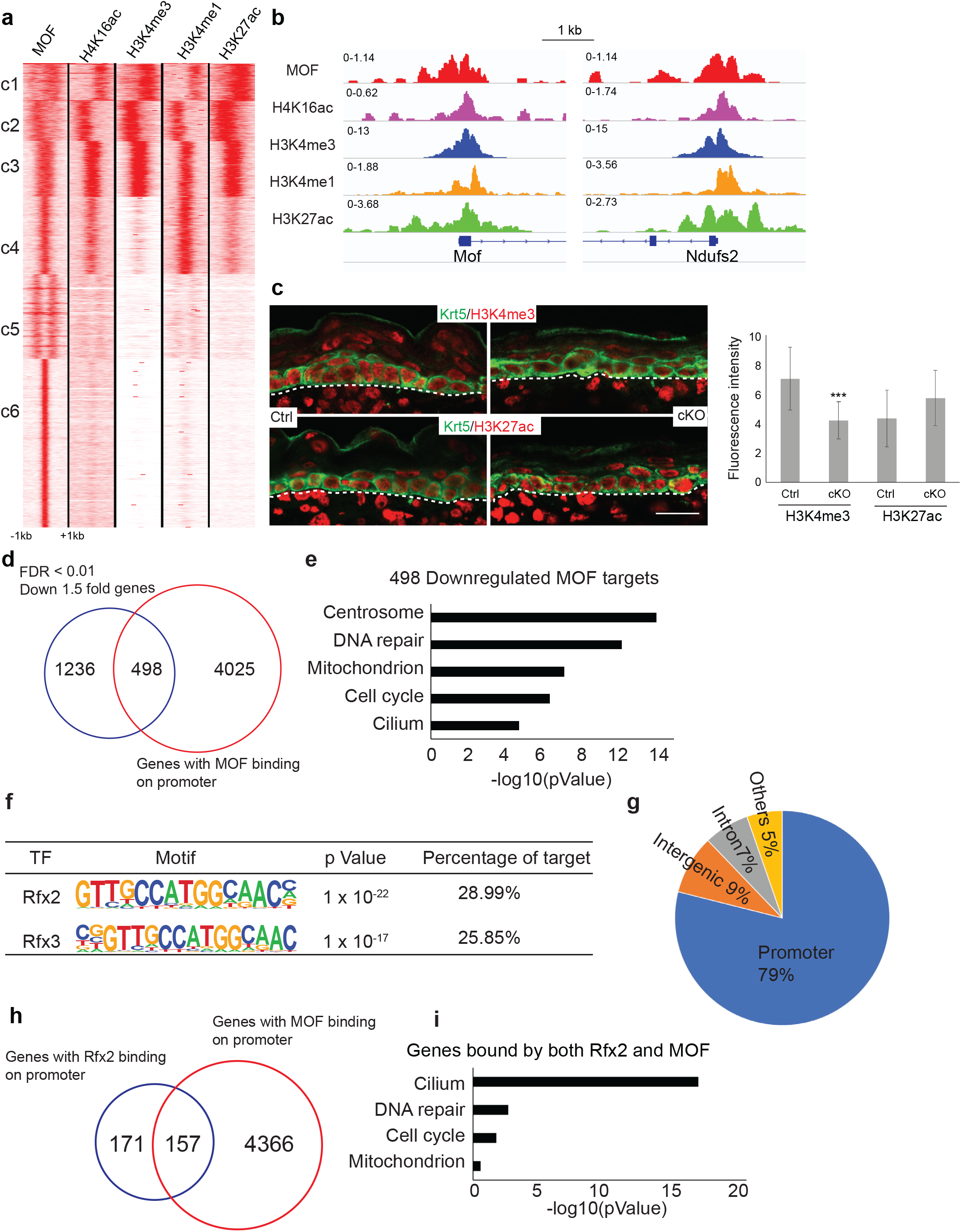
MOF regulates gene expression preferentially through promoters. **a**, *K*-means clustering of MOF ChIPseq, H4K16Ac, H3K4me3, H3K4me1 and H3K27Ac Cut&Run data for P0 epidermal keratinocytes. MOF ChIPseq peaks were used as genome coordinate, ± 1 kb from peak center. **b**, Example Integrative Genomics Viewer (IGV) browser view of tracks for two gene promoter regions: Mof and Ndufs2. **c**, Reduced promoter mark H3K4me3 and non-changed enhancer mark H3K27ac in cKO. White dashed lines mark epidermal-dermal boundary. **d**, Venn diagram of down-regulated genes in cKO with genes that have MOF binding on their promoter. **e**, GO terms for genes that not only down-regulated but also have MOF binding on their promoter, which we define as MOF targets. **f**, Motif enriched on down-regulated ciliary gene promoters. **g**, Peak distribution of Rfx2 Cut&Run. **h**, Venn diagram of genes with Rfx2 binding on their promoters and genes with MOF binding on their promoters. **i**, GO terms for genes with both Rfx2 and MOF binding on their promoter.

Because MOF has been shown to interact with MLL1, a H3K4 methyltransferase, and the interaction leads to coordination between H4K16ac and the methylation marks of H3K4^2^, we stained for H3K4me3 and H3K27ac, which mark active promoters^35^ and active enhancers^36,37^, respectively, in control and MOF cKO skin. Interestingly, H3K4me3 signals were specifically downregulated in MOF cKO epithelial cells but not in adjacent dermal cells (Fig. 5c). In contrast, H3K27ac signals were unchanged in both epithelial and dermal cells of MOF cKO (Fig. 5c). This result suggests that MOF and H4K16ac globally coordinate with H3K4me3 in epithelial cells, as one may expect.

Because genes that were downregulated upon MOF deletion showed a strong enrichment for MOF binding on their promoters, we next focused on these functional targets of MOF, which were defined as genes not only bound by MOF at their promoters but also downregulated in cKO (Fig. 5d). Indeed, genes associated with centrosome, DNA repair, mitochondrion, cell cycle and cilium are *bona fide* MOF direct targets (Fig. 5e).

To determine if any TFs coordinate with MOF and H4K16ac at gene promoters, we performed TF motif search at the promoter of downregulated genes in MOF cKO. Interestingly, RFX2 (Regulatory Factor binding to the X-box) and RFX3 motifs were identified as the most highly enriched motifs (Fig. 5f). Notably, RFX factors belong to an evolutionarily conserved family of TFs and play important roles in governing ciliary gene expression in *C*.*elegans*, mouse and human^38–40^. We next performed RFX2 Cut&Run to determine RFX2 footprint globally in epithelial cells of the skin (Extended Data fig. 4b). Interestingly, 79% of RFX2 bound regions (266 peaks) were on gene promoters (Fig. 5g). Among them, 48% (157 genes) of RFX2 bound promoters, including many ciliary genes such as *Ift74* and *Wdr34* (Extended Data fig. 4c), overlapped with MOF bound promoters (Fig. 5h). We noticed that the number of RFX2 bound gene promoters (328) were higher than the number of RFX2 bound peaks (266). This was caused by the head-to-head configuration of some RFX2 and MOF bound promoters in the genome, many of which are ciliary and mitochondrial genes and downregulated in MOF cKO (Extended Data fig. 4d-e). GO term analysis of these 157 genes revealed that cilium, DNA repair, cell cycle and mitochondrion were the most highly enriched gene categories co-regulated by RFX2 and MOF at the promoter (Fig. 5i). In contrast to the strong preference to promoters by RFX2, only 16% of regions bound by ΔNp63, a master TF of epithelial cell fate specification^27^, were on gene promoters whereas 80% of ΔNp63 bound regions were located in introns or intergenic regions (Extended Data fig. 4f). Furthermore, there was no enrichment for ΔNp63 regulated targets in either upregulated or downregulated genes in MOF cKO (Extended Data fig. 4g). Taken together, these results uncover the strong functional association and specificity of MOF and transcription factor RFX2 at gene promoters.

### MOF is required for maintaining chromatin accessibility at promoters

Because H4K16ac plays a critical role in chromatin decompaction^8,22^, we next performed scATAC-seq in control and MOF cKO cells isolated from E16.5 skin and determined the differential changes of chromatin accessibility at promoters and enhancers responding to the loss of MOF and H4K16ac. Overall, MOF deletion strongly altered the open chromatin landscape of the epithelial cells, slightly perturbed the dermal fibroblasts but largely spared other cell types in the skin, judging by the clustering patterns (Extended Data fig. 5a-b).

We next examined the open chromatin status of transcription start site (TSS). In epithelial cell clusters, the TSS enrichment score, calculated by ArchR^41^, was significantly reduced at MOF bound promoters in MOF cKO (Fig. 6a). As control, the TSS enrichment score remained unchanged for the dermal cells (Extended Data fig. 5c). Importantly, the normalized insertion score, quantifying the probability of Tn5 insertion and the openness of chromatin^41^, was significantly reduced for MOF targeted promoters but not at epithelial lineage gene promoters that are not bound by MOF (see Supplemental Table 2) or at the promoters of dermal cells (Fig. 6b-c and Extended Data fig. 5d). Furthermore, the normalized insertion score was significantly higher for MOF targeted promoters than that of the epithelial lineage genes (compare Fig. 6b and 6c). Collectively, these data demonstrate a highly opened chromatin state of MOF-bound promoters and provide experimental evidence for the requirement of MOF and H4K16ac for chromatin accessibility at targeted promoters.

**Fig. 6.**
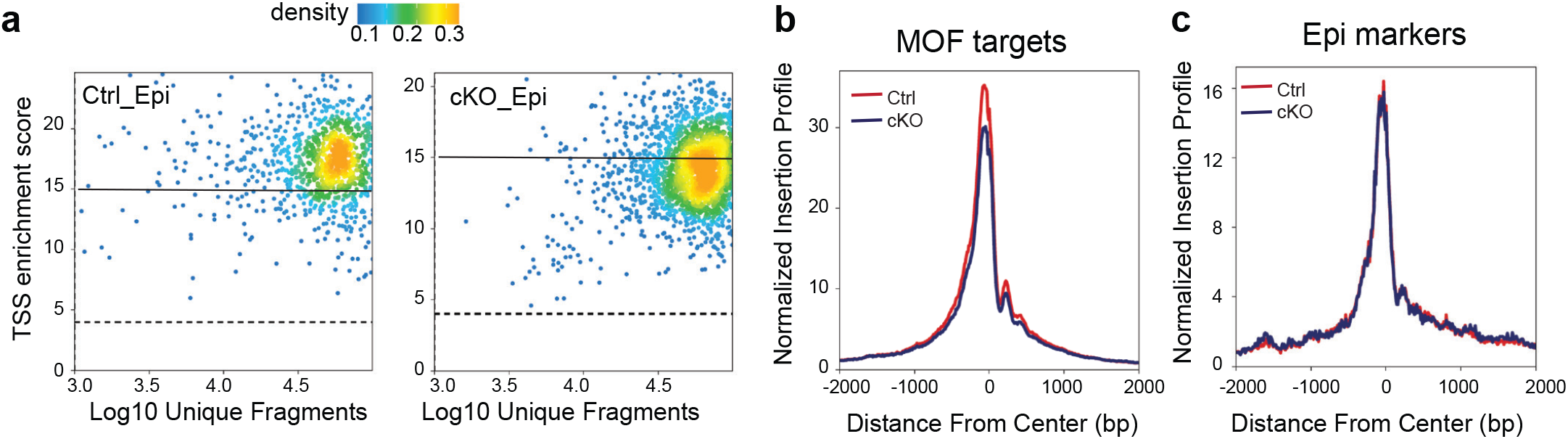
MOF is required for maintaining chromatin accessibility at promoters. **a**, Reduced TSS enrichment score in cKO epithelial cells. Ctrl_Epi, control epithelial cells, cKO_Epi, MOF cKO epithelial cells. **b**, Reduced normalized Tn5 insertion profile for MOF target genes. **c**, Similar normalized Tn5 insertion profile for epithelial marker genes.

### Genetic deletion of QPC recapitulates skin defects of MOF cKO

Having demonstrated that MOF-regulated gene expression program is important for mitochondria and cilia, we next sought to determine how compromised organelle functions cause the defects in MOF cKO. We noticed that nuclear encoded genes of the ETC, including the essential subunits of Complex I (*Ndufv1, Ndufb8, Ndufs2*), Complex II (*Sdha, Sdhb*), Complex III (*Uqcrc1, Uqcrc2*) and mitochondrial ATP synthase (*Atp5a1, Atp5b*), were downregulated in MOF cKO. These results prompted us to examine whether defects in MOF cKO were mediated, in part, by the compromised ETC. Because genetic deletion of ETC complex III component *Uqcrq* (QPC) disrupts the entire ETC^42^, we used *Krt14-Cre* to delete *Uqcrq* (QPC) and examined the phenotypes. Strikingly, QPC cKO mice died within hours after birth, mimicking MOF cKO. Although QPC cKO mice were similar in size comparing to WT or het control, their skin was thin and transparent, indicative of epidermal defects (Fig. 7a). Morphological analysis confirmed the thin epidermis and stunted HF growth such that more HFs failed to grow downward into the dermis (Fig. 7b and Extended Data fig. 6a). IF signals of KRT1, a spinous layer marker, and Lor, a granular layer marker, showed significantly reduced thickness of the epidermis and compromised epidermal differentiation (Fig. 7c). Distinct from MOF cKO skin, however, the BM was intact with the proper signals of β4 integrin (Extended Data fig. 6b) and epidermal proliferation showed no change (Extended Data fig. 6c). These data suggest that the epithelial defects of QPC cKO skin were manifested in epidermal differentiation and HF growth.

**Fig. 7.**
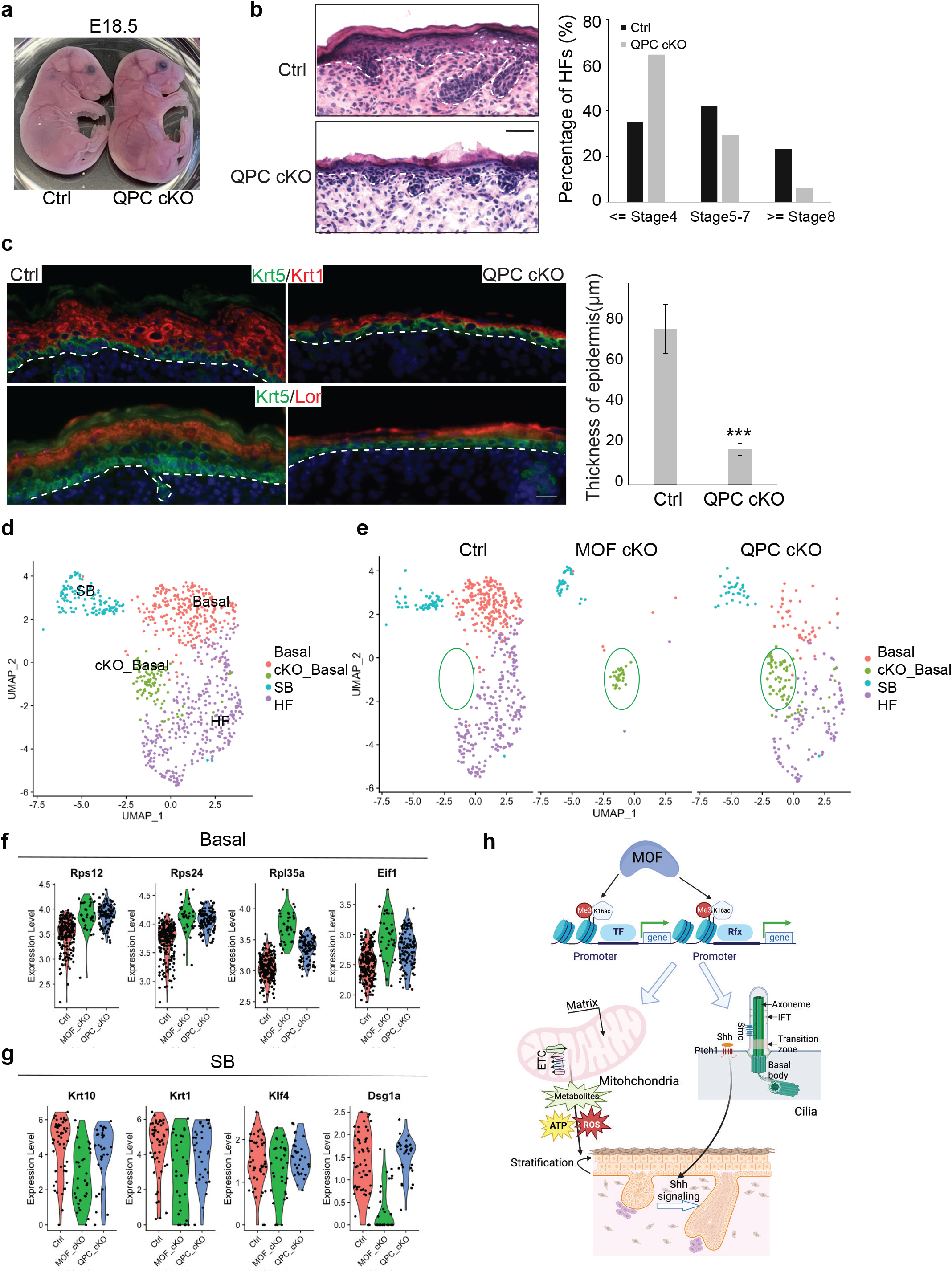
Genetic deletion of QPC recapitulates skin defects of MOF cKO. **a**, Relatively transparent appearance of QPC/Krt14 Cre cKO at E18.5. **b**, HE staining showing thinner epidermis and lacking of advanced stage hair follicles (HF). **c**, Differentiation defects and thinner epidermis indicated by Krt1 and Loricrin (Lor) staining. **d**, Integration and clustering of control and MOF cKO epithelial cells together with control and QPC cKO epithelial cells. **e**, Split view of control, MOF cKO and QPC cKO epithelial cells, as clustered in **d. f**, Examples of commonly up-regulated genes, related to protein translation, in MOF cKO and QPC cKO basal progenitor cells. **g**, Examples of commonly down-regulated, differentiation related genes in MOF cKO and QPC cKO suprabasal cells. **h**, Schematic illustration of MOF/H4K16ac-regulated gene expression controlling epidermal differentiation and hair growth through mitochondria and primary cilia. White dashed lines mark epidermal-dermal boundary. *P* value was calculated by unpaired two-tailed Student’s *t* test (**c**), ***, *P* < 0.001. Scale bar, 50 μm (**b)**, 20 μm (**c**).

To compare defective gene expression caused by the deletion of QPC to that of MOF cKO, we performed scRNA-seq of QPC cKO skin (Extended Data fig. 6d-e). Overall, the basal epithelial cell population, which self-renews and gives rise to differentiated suprabasal cells, were reduced in QPC cKO (Fig. 7d-e). In comparison, the basal cell population was largely depleted in MOF cKO (Fig. 7e). Interestingly, a unique basal epithelial cell population was detected in both MOF and QPC cKO but not in control skin (Fig. 7e), indicating similar changes of transcriptome and cellular state in the basal cells of MOF and QPC cKO. Closer inspection revealed that this basal epithelial cell population had highly elevated gene expression in ribosomal genes and translation (Fig. 7f), revealing a similar response of epidermal cells to the deletion of MOF and QPC. In addition, the basal cells in both MOF and QPC cKO had reduced AP1 TF expression, including *Fos, Jun, Junb* and *Atf3* (Extended Data fig. 6f), which play important roles for epidermal differentiation^27,43,44^. In suprabasal cells, the commonly downregulated genes were enriched in differentiation and keratin genes, such as *Krt10, Krt1, Klf4* and *Dsg1a* (Fig. 7g), and oxidoreductase activity, such as *Aldh2, Mdh1, Sdhb* and *Dhrs1* (Extended Data fig. 6g). Together, these data provide genetic and molecular evidence that MOF-controlled epidermal differentiation is mediated by its regulation of the ETC.

## Discussion

In this study, we used genetic mouse models and genomic tools to investigate the function and mechanism of MOF and H4K16ac in embryonic skin development. Leveraging spatiotemporally well-defined phenotypical and genomic analysis, we have uncovered specific functions of MOF-regulated transcriptome in governing epithelial cell adhesion, cell cycle, epidermal differentiation and HF growth (Fig. 7h). Importantly, genetic deletion of *Kansl1*, an essential component of NSL, leaves H4K16ac largely intact and causes minimal disruption of embryonic skin development, in sharp contrast to MOF cKO. Because previous studies have demonstrated that NSL has a broader substrate specificity and mediates the acetylation of non-histone proteins such as p53 and LSD1 as well as intramitochondrial functions of MOF^1,13,15,18^, these results strongly argue that histone acetyltransferase activities of MOF are essential for embryonic skin development. Furthermore, biochemical studies have demonstrated that the acetylation activity of MSL is largely restricted to H4K16 whereas NSL has a broader specificity to H4K5 and H4K8^16^. A recent study, using human acute myeloid leukemia cell line THP-1, suggested that the NSL complex is essential for cell survival and transcription initiation, likely through H4K5ac and H4K8ac but not H4K16ac^20^.

We note that cultured cells are usually selected against cell cycle defects, which could mask other functions of MOF and H4K16ac. In contrast, epidermal proliferation and differentiation are spatiotemporally distinct processes^45^, which permits our study of MOF and H4K16ac during skin development despite the strong defect in cell cycle in the basal cells. Collectively, our results suggest that MOF-mediated H4K16ac is essential for mammalian skin development and demonstrate epithelial cells of the skin as a model system to examine physiological function of H4K16ac *in vivo*.

In search for the complete set of MOF targeted genes in the skin, we performed both scRNAseq and bulk RNAseq. Surprisingly, we have identified ciliary genes, which escaped the detection in scRNAseq, and mitochondrial genes, which were detected in both single-cell and bulk RNAseq, as the major classes of MOF targets. Interestingly, ciliary genes are regulated by the conserved RFX TF family^38–40^, and the promoter regions of MOF-regulated ciliary genes are highly enriched for the motifs of RFX2 and RFX3. This serendipitous finding led us to determine that MOF-regulated promoters, many of which are co-bound by RFX2, have significantly more accessible chromatin than those of epithelial lineage genes, supporting the notion that H4K16ac regulates decompaction of the chromatin. Interestingly, RFX2 preferentially binds to promoters, in sharp contrast to ΔNp63, which prefers distal enhancers. Thus, these results nominate RFX2 as a new member of the class of TFs with a strong preference to gene promoters^46^. The coordination between RFX2 and MOF to regulate ciliary and mitochondrial genes provides an example for the interplay between TFs and histone H4 acetylation to maintain important cellular functions (Fig. 7h). Notably, a recent study of the NuA4 acetyltransferase complex in yeast provides a structural basis for the dual function of H4 acetylation in regulating chromatin packaging and transcription activation^47^. Thus, we speculate that MOF-mediated H4K16ac is an intrinsic mechanism to link chromatin decompaction and transcription activation.

We have identified mitochondria and cilia as two important organelles that are regulated by MOF. The function of primary cilia during skin development has been extensively studied recently. Notably, their function in epidermal differentiation^34^ and downgrowth of hair follicles^48,49^, through the control of Shh signaling, was reminiscent to defects observed in MOF cKO skin. In contrast, the role of mitochondria, in particular the specific function of the ETC, during embryonic skin development is unknown. Furthermore, whether MOF regulates mitochondrial function through direct intramitochondrial mechanisms or nuclear mechanisms is still under active debate^50^. Interestingly, genetic ablation of Complex III of the ETC, which disrupts the entire ETC^42^, has recapitulated the severe defects in epidermal differentiation and hair growth. Notably, previous studies have implicated MOF in controlling mitochondrial functions by activating fatty acid oxidation pathway^3^ or regulating mitochondrial transcription and respiration through a subset of the NSL complex within the mitochondria^1^, respectively. Our results have identified the ETC as another major mitochondrial function regulated by this versatile layer of gene regulation. Of note, genetic deletion of *Tfam* (transcription factor A, mitochondrial), a DNA-binding protein essential for transcriptional activation of mtDNA^51^, caused defects in epidermal differentiation and HF development but the animals survived more than 2 weeks after birth^52^, which is less severe than MOF and QPC cKO. Therefore, MOF-mediated nuclear mitochondrial gene expression plays a major role in the skin. Future investigation will be needed to elucidate how the ETC functions, including the production of ATP, ROS or other metabolites, promote epidermal differentiation and hair growth in a MOF-dependent manner.

In conclusion, this study provides genetic, genomic and molecular evidence for MOF-mediated regulation of mitochondria and primary cilia and demonstrates the coordination between RFX2 and MOF to govern gene expression in mammals. This research now paves the way to probe more deeply into the intricate network of mitochondria, cilia and MOF-controlled epigenetic regulation in development, homeostasis and disease.

### Methods Mice

All experiments were carried out following IACUC-approved protocols and guidelines at CU Boulder and Northwestern, respectively. Mice were bred and housed according to guidelines of the IACUC in a pathogen-free facility at University of Colorado at Boulder and at Northwestern University Feinberg School of Medicine.

MOF fl/fl line was generated as described previously^5^. Kansl1 fl/fl line was generated from a conditional ready, targeted ESC strain obtained from the European Mouse Mutant Archive (EMMA). In this conditional allele, exon 3 (out of total 14 exons) is targeted for Cre-mediated recombination and deletion. Other mouse lines used include: Krt14-Cre (E. Fuchs, Rockefeller University), Krt14-H2BGFP (E. Fuchs, Rockefeller University).

### Immunofluorescence and H&E staining

OCT-embedded embryos or dorsal skin tissues were sectioned to 10 μm, fixed in 4% PFA for 10 min at room temperature (RT), permeabilized with 0.1% Triton X-100 for 10 min at RT. For BrdU staining, samples were treated with 24 U/mL DNase in DNase Buffer (30 mM Tris-Cl, pH8.1, 0.5 mM β-Mercaptoethanol, 30 mM MgCl_2_) for 1 hr at 37°C before proceeding to the next step. When staining with mouse derived antibodies, mouse-on-mouse basic kit (BMK-2202, Vector Laboratories) was used. Otherwise, blocking was performed with 5% normal serum of the same species that the secondary antibody was raised in. Sections were incubated with primary antibodies overnight at 4°C. Next day, sections were washed three time for 5 min each, then incubated with secondary antibodies for 2 hrs at RT. Nuclei were stained with Hoechst33342 (1: 5,000, Invitrogen). Slides were then mounted with VECTASHIELD antifade mounting medium (Vector Laboratories, H-1000). Imaging was performed either on a Leica DM5500B microscope with an attached Hamamatsu C10600-10B camera and MetaMorph software, or a Nikon A1 confocal microscope. Primary antibodies and dilution used in this study include H4K16ac (Sigma-Aldich, #07-329, 1:2000), Krt5 (Covance, #SIG-3475, 1:2000), Krt1 (Covance, #PRB-165P, 1:2000), Loricrin (The Rockefeller University, Gift from E. Fuchs), β4 integrin (BD Biosciences, #553745, 1:200), Col17 (Abcam, #ab184996, 1:500), Pericentrin (Covance, #PRB-432C, 1:200), E-Cadherin (The Rockefeller University, Gift from E. Fuchs, 1:200), α-Catenin (Cell Signaling Technology, #3236, 1:1000), Active-Caspase3 (R&D Systems, #AF835, 1:1000), Sox9 (Millipore, #AB5535, 1:500), Ki67 (Abcam, #ab15580, 1:500), Lef1 (Cell Signaling Technology, #2230, 1:500), P63 (Cell Signaling Technology, #4892, 1:200), BrdU (Abcam, #ab6326, 1:500), Krt6 (Covance, #PRB-169P, 1:500), H3K4me3 (Cell Signaling Technology, #9751, 1:200), H3K27ac (Cell Signaling Technology, #8173, 1:200). All secondary antibodies were from Invitrogen/Molecular Probes, used at 1:2000.

Wholemount staining of Arl13b (NeuroMab, #AB_11000053, 1:10) in Krt14-H2BGFP^+^ samples was used for cilia detection. The protocol was adapted from a previous report^53^. Briefly, freshly dissected Krt14-H2BGFP^+^ E16.5 control and MOF cKO dorsal skin samples were fixed in 4% PFA, blocked in PB Buffer (0.5% skim milk powder, 0.25% fish skin gelatin, 0.5% Triton X-100 in 1x PBS) for 1 hr at RT. One drop of M.O.M blocking reagent from the mouse-on-mouse basic kit (BMK-2202, Vector Laboratories) was added per 1 mL PB buffer during the blocking since Arl13b is a mouse antibody. Arl13b was diluted in PB buffer and the samples were incubated with primary antibody overnight. Next day, the samples were washed twice at 2∼3 hrs each time, and then incubated with 1:1000 secondary antibody and 1:5000 Hoechst33342 overnight. The third day, samples were changed to PBS, without thorough washes, mounted onto slides with VECTASHIELD antifade mounting medium. Imaging was performed on a Nikon A1 confocal microscope.

For H&E staining, sections were fixed in 4% PFA for 10 min at RT, then dipped in hematoxylin for 3 min, eosin for 6 sec. After going through a series of dehydration steps, sections were mounted with Permount mounting medium (Fisher Scientific, SP15-500). Imaging was performed on a Leica DM5500B microscope with a Leica camera.

### Flow cytometry

For E16.5 sorting, total dorsal skin was mined into small pieces and incubated with 0.1% collagenase (Worthington, LS004188) for 30 min at 37°C. After incubation, PBS was used to dilute the collagenase, then pipet to dissociate. Tissues were pelleted by centrifuge and then subject to fresh Trypsin digestion for 5 min at 37°C. PBS supplemented with 5% chelated FBS was used to neutralize Trypsin and cells were filtered through 40-μm cell strainer. DAPI-positive dead cells were excluded and epithelial cells were enriched by selecting Krt14-H2BGFP+ cells. Flow cytometry was performed on the MoFlo XDP machine (Beckman Coulter).

For P0.5 newborn epidermal cell isolation, dorsal skin was first incubated with 2.5 U/mL Dispase to separate epidermis from underlying dermis. Then epidermis was floated on Trypsin, incubating for 5 min at 37°C. Single cell suspension was generated by pipetting and filtering through 40-μm cell strainer and used for downstream experiments.

### Electron Microscopy

E16.5 control and MOF cKO dorsal skin samples were fixed in EM fixation solution (2% Glutaraldehyde, 4% PFA, 50 mM Na Cacodylate, 2 mM Calcium Chloride) in a glass vial, and then bring to the CU Boulder EM Service Core Facility for further processing. Imaging was performed using the FEI Tecnai F30, 300kV FEG-TEM system.

### shRNA infection

pGIPZ vector clone expressing control or Msl1 shRNA were obtained from Northwestern SBDRC Gene Editing, Transduction and Nanotechnology (GET iN) core. Lentivirus were packed in HEK293T cells. Lentivirus supernatant was added to wildtype mouse keratinocytes in 6-well plate overnight to infect. 24 hrs after infection, 1 μg/mL puromycin was added to the cells to select positively infected cells. 72 hrs after selection, cells were collected for qPCR or plated onto cover glass for immunofluorescence staining.

### Reverse transcription and real-time PCR

Reverse transcriptions were performed using SuperScript III First-Strand Synthesis SuperMix (Invitrogen, 18080-400). Real-time PCR reagents were from Bio-Rad. Reactions were performed according to the manufacturer’s manual on a CFX384 real-time system (Bio-Rad). Differences between samples and controls were calculated using the 2 ^-ΔΔC(t)^ method.

### RNAseq library construction and analysis

Total RNA was isolated from flow cytometry enriched E16.5 control and MOF cKO epithelial cells using TRIzol (Invitrogen), and RNA quality was assessed with Agilent 2100 bioanalyer. 1 μg total RNA was used for RNAseq library construction using NEBNext Ultra Directional RNA Library Prep Kit for Illumina (NEB, E7420L) per the manufacturer’s protocol.

For Kansl1 RNAseq, which was performed at P0.5, the epidermis was separated from dermis after Dispase treatment, and then directly floated on 1 mL TRIzol reagent, incubated at RT for 15 min. The lysate was collected for RNA purification and library construction as described above.

RNAseq reads (150 nt, paired-end) were aligned to the mouse genome (NCBI37/mm10) using HISAT2 (version 2.1.0)^54^. Expression of each gene was calculated from the alignment Bam file by HTSeq-count^55^. Differentially expressed genes were determined using R package DESeq2^56^ with padj < 0.01, 1.5x fold-change as cutoff. Gene ontology analysis for changed genes was performed using DAVID Bioinformatics Resources 6.8^57^. GSEA^32^ analysis was performed using fold change values (MOF cKO *vs*. control) as the expression dataset, and gene sets were downloaded from GSEA website for ciliary gene set and mitochondrial gene set or curated for p63 targets from a previous publication^27^.

### ChIPseq and Cut&Run

All ChIPseq and Cut&Run experiments were performed using newborn total epidermal cells. For MOF ChIPseq, epidermal single cell suspension was subject to sonication, then pulled down using MOF antibody (Bethyl, #A300-992A). Cut&Run was conducted following this protocol: https://protocols.io/view/cut-amp-run-targeted-in-situ-genome-wide-profiling-zcpf2vn.html. Cut&Run antibodies used in this study includes H4K16ac (Sigma-Aldich, #07-329), H3K4me3 (Cell Signaling Technology, #9751), H3K4me1 (Cell Signaling Technology, #9723), H3K27ac (Cell Signaling Technology, #8173), Rfx2 (Sigma-Aldrich, #HPA048969). All used at 1:100. The libraries were constructed using NEBNext^®^ Ultra™ II DNA Library Prep Kit for Illumina^®^.

Single-end ChIPseq and Paired-end Cut&Run reads were aligned to the mouse genome (NCBI37/mm10) using Bowtie2^58^ with option ‘‘--very-sensitive-local’’. Peak calling was performed on each individual sample by MACS2^59^. Parameter ‘‘-BAMPE’’ was used for Rfx2 peak calling.

*K*-means clustering was performed using seqMINER (version 1.3.4)^60^. First, mapped Bam files were converted to Bed files using bedtools, then 10 million (MOF ChIPseq and H4K16ac, H3k4me1, H3K27ac Cut&Run) or 2 million (H3K4me3) mapped reads were randomly sampled from each library and used as input. MOF ChIPseq peaks were used as genome coordinate. Signals were calculated in a 2 kb region (±1,000 bp) surrounding the center of the peak with 25 nt bin.

Enriched motif search was performed using Homer^61^ *findMotifGenome*.*pl*. Peak distribution in the genome is annotated using Homer *annotatePeaks*.*pl* function.

### single-cell RNAseq library generation

Single cell suspensions from two pairs of E16.5 control and MOF cKO total dorsal skin were used as input for single cell RNAseq. The libraries were constructed following 10x Genomics Chromium Single Cell 3’ Reagent Kits v2 User Guide (PN-120237). The libraries were sequenced on the Illumina NovaSeq 6000 platform to achieve an average of approximately 50,000 reads per cell.

For QPC scRNAseq, E17.5 control and QPC cKO total dorsal skin were used as input. The libraries were constructed following 10x Genomics Chromium Single Cell 3’ Reagent Kits v3 User Guide (PN-1000075). The libraries were sequenced on the Illumina NovaSeq 6000 platform to achieve an average of approximately 50,000 reads per cell.

### single-cell RNAseq analysis

The 10x Genomics Cellranger Single-Cell Software Suite was used to perform barcode processing and single-cell 3′ gene counting. Barcodes, features and matrix files were loaded into Seurat 4.0^62^ for downstream analysis. Low quality and dead cells were excluded from analysis using nFeature_RNA (>1000 and <6,000) and mitochondrial percentage (<10%). In addition, cell cycle regression was used to regress out addition variation from cell cycle genes. After UMAP dimension reduction and clustering, cluster markers were used to identify distinct cell populations. Genes with log2FC more than 0.5 were used for GO term analysis.

To analyze epithelial cell in higher resolution, we subset epithelial cells and then re-ran the analysis pipeline.

To calculate nuclear mitochondrial score, a list of 1,158 mitochondrial genes downloaded from GSEA were used to calculate a module score using *AddModuleScore* function of Seurat.

To make the cell number comparable, 600 epithelial cells were randomly subset from QPC control and cKO samples, and then integrated with MOF epithelial cells using Seurat 4.0 for further analysis.

### single-cell ATACseq library generation

For scATACseq, E16.5 control and MOF cKO total dorsal skins were subject to flow cytometry to sort for Krt14-H2BGFP^+^ epithelial cells and Krt14-H2BGFP^-^ non-epithelial cells, then the GFP^+^ cells were mixed with GFP^-^ cells at 2:1 ratio to enrich for epithelial cells. The libraries were prepared using the 10x Genomics Chromium Single Cell ATAC Library & Gel Bead kit (PN-1000110). The libraries were sequenced on the Illumina NovaSeq 6000 platform to achieve an average of approximately 50,000 paired-end reads per nucleus.

### single-cell ATACseq analysis

Raw sequencing reads were aligned using cellranger-atac (version 2.0.0) with corresponding reference genome (refdata-cellranger-arc-mm10-2020-A-2.0.0) downloaded from 10x Genomics website. The generated *filtered_peak_bc_matrix* and *fragment* files were used for further analysis.

R package ArchR^41^ was used for cell clustering, TSS enrichment score calculation and Tn5 insertion profile plotting. MOF targets were identified as genes that have MOF binding on their promoter and down-regulated in MOF cKO. Epithelial markers were identified from scRNAseq.

### Statistics and study design

RNAseq and scRNAseq for control and MOF cKO were performed with two pairs of samples. scATAC-seq was performed with one pair of control and MOF cKO samples. All the experiments were designed such that there were always littermate controls. All statistical tests were performed using unpaired two-tailed Student’s *t* test. No statistical methods were used to predetermine sample size. Experiments were not randomized, and investigators were not blinded to allocation during experiments or outcome assessment.

## Data availability

All sequencing data were deposited to NCBI/GEO Super Series under accession number GSE214441.

## Acknowledgements

We thank E. Fuchs (Rockefeller University, HHMI) for Krt14-H2bGFP and Krt14-Cre mice, J. Wallingford (University of Texas at Austin) for suggestions regarding to ciliary studies, and all members of the Yi laboratory for suggestions. This work was supported by National Institute of Health Grant R01AR071435, R01AR081103 and R01HD107841 (RY).

## Contributions

D.W. performed most experiments and analyses with assistance from H.L.; H.L. performed ChIP-seq and Cut&Run profiling; N.S.C. provided the QPC mouse model and advised the mitochondrial study; Y.D. provided the MOF mouse model and advised the MOF/H4K16ac study; R.Y. conceived and supervised the study and wrote the manuscript together with D.W. All authors participated in the manuscript preparation.

## Conflict of interests

The authors declare no conflict of interests.

## Figure Legend

**Extended Data Fig. 1** | **Kansl1 knockout strategy and phenotypical analysis. a**, Schematic illustration of the generation of Kansl1 conditional allele. **b**, Representative genotyping result by genomic DNA PCR. Homo, homozygous, Kansl1 fl/fl; Het, heterozygous, Kansl fl/+; WT, Kansl1 +/+. **c**, RNAseq track showing deletion of exon3 in Kansl1 cKO. **d**, Increased apoptosis in Kansl1 cKO shown by active-caspase 3 (Ac-Caspase3) staining. **e**, Immunofluorescence staining showing non-discernible difference between control and Kansl1 cKO in basement membrane deposition (β4 integrin), differentiation and epidermal thickness (Krt1), hair follicle stem cell specification (Sox9) and proliferation (Ki67). **f**, Volcano plot of differentially expressed genes in Kansl1 cKO compared with control. Red dots represent genes showing fold change > 1.5 and FDR < 0.01. Ctrl, control. White dashed lines indicate epidermal-dermal boundary (**d** and **e**), *P* value was calculated by unpaired two-tailed Student’s *t* test (**d**), **, *P* < 0.01. Scale bar, 20 μm (**d** and **e**).

**Extended Data Fig. 2** | **Phenotypical analysis of MOF cKO. a**, Confirmation of epithelial fate specification in cKO by epithelial master transcription factor P63 staining. **b**, Proliferation examination by BrdU staining. **c**, Apoptosis detected by active-Caspase 3 (Ac-Casp3). **d**, Stress response indicated by Krt6 staining in the epidermal differentiated layer. **e**, Specification of hair follicle stem cells in the few hair follicles detected in cKO as shown by Sox9 staining. **f**, Hair follicle specification demonstrated by Lef1 staining. Ctrl, control. White dashed lines indicate epidermal-dermal boundary (**a**-**f**), *P* values were calculated by unpaired two-tailed Student’s *t* test (**b** and **c**), ***, *P* < 0.001. Scale bar, 20 μm (**a**-**f**).

**Extended Data Fig. 3** | **Single cell RNAseq analysis. a**, UMAP clustering of all the cells detected in scRNAseq, colored by genotype. **b**, UMAP clustering of epithelial cells, colored by genotype. **c**, Feature plot of genes that were used for identifying each epithelial cluster. **d**, Violin plots showing reduced basement membrane genes detected by scRNAseq. **e**, GO terms of down-regulated genes in cKO *vs* control suprabasal comparison. **f**, Example of down-regulated mitochondrial matrix genes.

**Extended Data Fig. 4** | **RNAseq and ChIPseq analysis. a**, Log2 fold change of down-regulated basement membrane genes detected by RNAseq. **b**, Top enriched motif in Rfx2 Cut&Run peaks. **c**, Examples of two ciliary genes, Ift174 and Wdr34, that have both MOF and Rfx2 binding on their promoters marked by H3K4me3. **d**, Examples of two ciliary genes, Ift122 and Tmem138, that share one promoter with another genes, have both MOF and Rfx2 binding on their promoters, and the two genes are both down-regulated. **e**, Examples of two mitochondrial genes Lyrm4 and Fars2 that share one promoter, another mitochondrial gene Sirt3 shares the same promoter as Psmd13. Both shared promoters have MOF and Rfx2 binding and all four genes are down-regulated in cKO. **f**, Distribution of P63 Cut&Run peaks in the genome. **g**, GSEA for P63 targets identified in a previous study^27^. Values in parentheses indicate gene expression fold change calculated from RNAseq. Arrows indicate the direction of gene transcription.

**Extended Data Fig. 5** | **scATACseq analysis. a** and **b**, UMAP clustering of scATACseq data, colored by sample (**a**) or by cluster (**b**). **c**, TSS enrichment score for dermal population. Ctrl_Derm, control_dermal cells; cKO_Derm, MOF cKO_dermal cells. **d**, Normalized Tn5 insertion profile for all genes in dermal cells.

**Extended Data Fig. 6** | **QPC cKO phenotypical analysis. a**, Reduced advanced stage hair follicles in QPC cKO as shown by Lef1 staining. **b**, Intact basement membrane as indicated by β4 integrin staining. **c**, Non-changed proliferation marked by similar proportion of Ki67^+^ cells. **d**, Nine cell types detected in E17 dorsal skin samples by scRNAseq. **e**, Feature plot of genes that were used for identifying each epithelial cluster. **f**, Examples of commonly down-regulated AP1 TFs in MOF cKO and QPC cKO basal progenitor cells. **g**, Examples of down-regulated oxidoreductase activity genes in both MOF cKO and QPC cKO suprabasal cells. Ctrl, control.

## Notes

### Competing Interest Statement

The authors have declared no competing interest.

## References

1. Chatterjee, A. et al. MOF Acetyl Transferase Regulates Transcription and Respiration in Mitochondria. Cell 167, 722–738.e23 (2016).

2. Dou, Y. et al. Physical association and coordinate function of the H3 K4 methyltransferase MLL1 and the H4 K16 acetyltransferase MOF. Cell 121, 873–885 (2005).

3. Khoa, L. T. P. et al. Histone Acetyltransferase MOF Blocks Acquisition of Quiescence in Ground-State ESCs through Activating Fatty Acid Oxidation. Cell Stem Cell 27, 441–458.e10 (2020).

4. Kind, J. et al. Genome-wide analysis reveals MOF as a key regulator of dosage compensation and gene expression in Drosophila. Cell 133, 813–828 (2008).

5. Li, X. et al. The histone acetyltransferase MOF is a key regulator of the embryonic stem cell core transcriptional network. Cell Stem Cell 11, 163–178 (2012).

6. Samata, M. et al. Intergenerationally Maintained Histone H4 Lysine 16 Acetylation Is Instructive for Future Gene Activation. Cell 182, 127–144.e23 (2020).

7. Dion, M. F., Altschuler, S. J., Wu, L. F. & Rando, O. J. Genomic characterization reveals a simple histone H4 acetylation code. Proc Natl Acad Sci U S A 102, 5501–5506 (2005).

8. Zhang, R., Erler, J. & Langowski, J. Histone Acetylation Regulates Chromatin Accessibility: Role of H4K16 in Inter-nucleosome Interaction. Biophysical Journal 112, 450–459 (2017).

9. Valencia-Sánchez, M. I. et al. Regulation of the Dot1 histone H3K79 methyltransferase by histone H4K16 acetylation. Science 371, eabc6663 (2021).

10. Zhang, Y. et al. Selective binding of the PHD6 finger of MLL4 to histone H4K16ac links MLL4 and MOF. Nat Commun 10, 2314 (2019).

11. Taipale, M. et al. hMOF histone acetyltransferase is required for histone H4 lysine 16 acetylation in mammalian cells. Mol Cell Biol 25, 6798–6810 (2005).

12. Hilfiker, A., Hilfiker-Kleiner, D., Pannuti, A. & Lucchesi, J. C. mof, a putative acetyl transferase gene related to the Tip60 and MOZ human genes and to the SAS genes of yeast, is required for dosage compensation in Drosophila. EMBO J 16, 2054–2060 (1997).

13. Li, X., Wu, L., Corsa, C. A. S., Kunkel, S. & Dou, Y. Two mammalian MOF complexes regulate transcription activation by distinct mechanisms. Mol Cell 36, 290–301 (2009).

14. Mendjan, S. et al. Nuclear pore components are involved in the transcriptional regulation of dosage compensation in Drosophila. Mol Cell 21, 811–823 (2006).

15. Sheikh, B. N., Guhathakurta, S. & Akhtar, A. The non-specific lethal (NSL) complex at the crossroads of transcriptional control and cellular homeostasis. EMBO Rep 20, e47630 (2019).

16. Cai, Y. et al. Subunit composition and substrate specificity of a MOF-containing histone acetyltransferase distinct from the male-specific lethal (MSL) complex. J Biol Chem 285, 4268–4272 (2010).

17. Shvedunova, M. & Akhtar, A. Modulation of cellular processes by histone and non-histone protein acetylation. Nat Rev Mol Cell Biol 23, 329–349 (2022).

18. Luo, H. et al. MOF Acetylates the Histone Demethylase LSD1 to Suppress Epithelial-to-Mesenchymal Transition. Cell Reports 15, 2665–2678 (2016).

19. Chelmicki, T. et al. MOF-associated complexes ensure stem cell identity and Xist repression. eLife 3, e02024 (2014).

20. Radzisheuskaya, A. et al. Complex-dependent histone acetyltransferase activity of KAT8 determines its role in transcription and cellular homeostasis. Molecular Cell 81, 1749–1765.e8 (2021).

21. Ravens, S. et al. Mof-associated complexes have overlapping and unique roles in regulating pluripotency in embryonic stem cells and during differentiation. eLife 3, e02104 (2014).

22. Shogren-Knaak, M. et al. Histone H4-K16 Acetylation Controls Chromatin Structure and Protein Interactions. Science 311, 844–847 (2006).

23. Smith, E. R. et al. A human protein complex homologous to the Drosophila MSL complex is responsible for the majority of histone H4 acetylation at lysine 16. Mol Cell Biol 25, 9175–9188 (2005).

24. Vasioukhin, V., Degenstein, L., Wise, B. & Fuchs, E. The magical touch: Genome targeting in epidermal stem cells induced by tamoxifen application to mouse skin. Proc. Natl. Acad. Sci. U.S.A. 96, 8551–8556 (1999).

25. Ravens, S. et al. Mof-associated complexes have overlapping and unique roles in regulating pluripotency in embryonic stem cells and during differentiation. eLife 3, e02104 (2014).

26. Lechler, T. & Fuchs, E. Asymmetric cell divisions promote stratification and differentiation of mammalian skin. Nature 437, 275–280 (2005).

27. Fan, X. et al. Single Cell and Open Chromatin Analysis Reveals Molecular Origin of Epidermal Cells of the Skin. Dev. Cell 47, 21–37.e5 (2018).

28. Zhou, P., Byrne, C., Jacobs, J. & Fuchs, E. Lymphoid enhancer factor 1 directs hair follicle patterning and epithelial cell fate. Genes Dev. 9, 700–713 (1995).

29. Gat, U., DasGupta, R., Degenstein, L. & Fuchs, E. De Novo hair follicle morphogenesis and hair tumors in mice expressing a truncated beta-catenin in skin. Cell 95, 605–614 (1998).

30. St-Jacques, B. et al. Sonic hedgehog signaling is essential for hair development. Curr. Biol. 8, 1058–1068 (1998).

31. Chiang, C. et al. Essential Role forSonic hedgehogduring Hair Follicle Morphogenesis. Developmental Biology 205, 1–9 (1999).

32. Subramanian, A. et al. Gene set enrichment analysis: A knowledge-based approach for interpreting genome-wide expression profiles. PNAS 102, 15545–15550 (2005).

33. Reiter, J. F. & Leroux, M. R. Genes and molecular pathways underpinning ciliopathies. Nat Rev Mol Cell Biol 18, 533–547 (2017).

34. Ezratty, E. J. et al. A role for the primary cilium in Notch signaling and epidermal differentiation during skin development. Cell 145, 1129–1141 (2011).

35. Santos-Rosa, H. et al. Active genes are tri-methylated at K4 of histone H3. Nature 419, 407–411 (2002).

36. Creyghton, M. P. et al. Histone H3K27ac separates active from poised enhancers and predicts developmental state. PNAS 107, 21931–21936 (2010).

37. Rada-Iglesias, A. et al. A unique chromatin signature uncovers early developmental enhancers in humans. Nature 470, 279–283 (2011).

38. Chung, M.-I. et al. RFX2 is broadly required for ciliogenesis during vertebrate development. Developmental Biology 363, 155–165 (2012).

39. Swoboda, P., Adler, H. T. & Thomas, J. H. The RFX-type transcription factor DAF-19 regulates sensory neuron cilium formation in C. elegans. Mol Cell 5, 411–421 (2000).

40. Choksi, S. P., Lauter, G., Swoboda, P. & Roy, S. Switching on cilia: transcriptional networks regulating ciliogenesis. Development 141, 1427–1441 (2014).

41. Granja, J. M. et al. ArchR is a scalable software package for integrative single-cell chromatin accessibility analysis. Nat Genet 53, 403–411 (2021).

42. Diebold, L. P. et al. Mitochondrial complex III is necessary for endothelial cell proliferation during angiogenesis. Nat Metab 1, 158–171 (2019).

43. Eckert, R. L., Crish, J. F., Banks, E. B. & Welter, J. F. The epidermis: genes on - genes off. J. Invest. Dermatol. 109, 501–509 (1997).

44. Rorke, E. A., Adhikary, G., Young, C. A., Roop, D. R. & Eckert, R. L. Suppressing AP1 factor signaling in the suprabasal epidermis produces a keratoderma phenotype. J. Invest. Dermatol. 135, 170–180 (2015).

45. Mesa, K. R. et al. Homeostatic Epidermal Stem Cell Self-Renewal Is Driven by Local Differentiation. Cell Stem Cell 23, 677–686.e4 (2018).

46. Bergman, D. T. et al. Compatibility rules of human enhancer and promoter sequences. Nature 607, 176–184 (2022).

47. Qu, K., Chen, K., Wang, H., Li, X. & Chen, Z. Structure of the NuA4 acetyltransferase complex bound to the nucleosome. Nature 610, 569–574 (2022).

48. Chen, J. et al. The ciliopathy gene Rpgrip1l is essential for hair follicle development. J Invest Dermatol 135, 701–709 (2015).

49. Yang, N. et al. Intraflagellar transport 27 is essential for hedgehog signaling but dispensable for ciliogenesis during hair follicle morphogenesis. Development 142, 2194–2202 (2015).

50. Rubalcava-Gracia, D., García-Villegas, R. & Larsson, N.-G. No role for nuclear transcription regulators in mammalian mitochondria? Molecular Cell (2022) doi:10.1016/j.molcel.2022.09.010.

51. Bonawitz, N. D., Clayton, D. A. & Shadel, G. S. Initiation and Beyond: Multiple Functions of the Human Mitochondrial Transcription Machinery. Molecular Cell 24, 813–825 (2006).

52. Hamanaka, R. B. et al. Mitochondrial reactive oxygen species promote epidermal differentiation and hair follicle development. Sci Signal 6, ra8 (2013).

53. Driskell, R. R., Giangreco, A., Jensen, K. B., Mulder, K. W. & Watt, F. M. Sox2-positive dermal papilla cells specify hair follicle type in mammalian epidermis. Development 136, 2815–2823 (2009).

54. Zhang, Y., Park, C., Bennett, C., Thornton, M. & Kim, D. Rapid and accurate alignment of nucleotide conversion sequencing reads with HISAT-3N. Genome Res (2021) doi:10.1101/gr.275193.120.

55. Anders, S., Pyl, P. T. & Huber, W. HTSeq--a Python framework to work with high-throughput sequencing data. Bioinformatics 31, 166–169 (2015).

56. Love, M. I., Huber, W. & Anders, S. Moderated estimation of fold change and dispersion for RNA-seq data with DESeq2. Genome Biol. 15, 550 (2014).

57. Huang, D. W., Sherman, B. T. & Lempicki, R. A. Systematic and integrative analysis of large gene lists using DAVID bioinformatics resources. Nat Protoc 4, 44–57 (2009).

58. Langmead, B. & Salzberg, S. L. Fast gapped-read alignment with Bowtie 2. Nat. Methods 9, 357–359 (2012).

59. Zhang, Y. et al. Model-based Analysis of ChIP-Seq (MACS). Genome Biology 9, R137 (2008).

60. Ye, T. et al. seqMINER: an integrated ChIP-seq data interpretation platform. Nucleic Acids Res. 39, e35 (2011).

61. Heinz, S. et al. Simple Combinations of Lineage-Determining Transcription Factors Prime cis-Regulatory Elements Required for Macrophage and B Cell Identities. Molecular Cell 38, 576–589 (2010).

62. Hao, Y. et al. Integrated analysis of multimodal single-cell data. Cell 184, 3573–3587.e29 (2021).

