## Supplementary Data for "MOF-mediated Histone H4 Lysine 16 Acetylation Governs Mitochondrial and Ciliary Functions By Controlling Gene Promoters"

### Extended Data Fig. 1

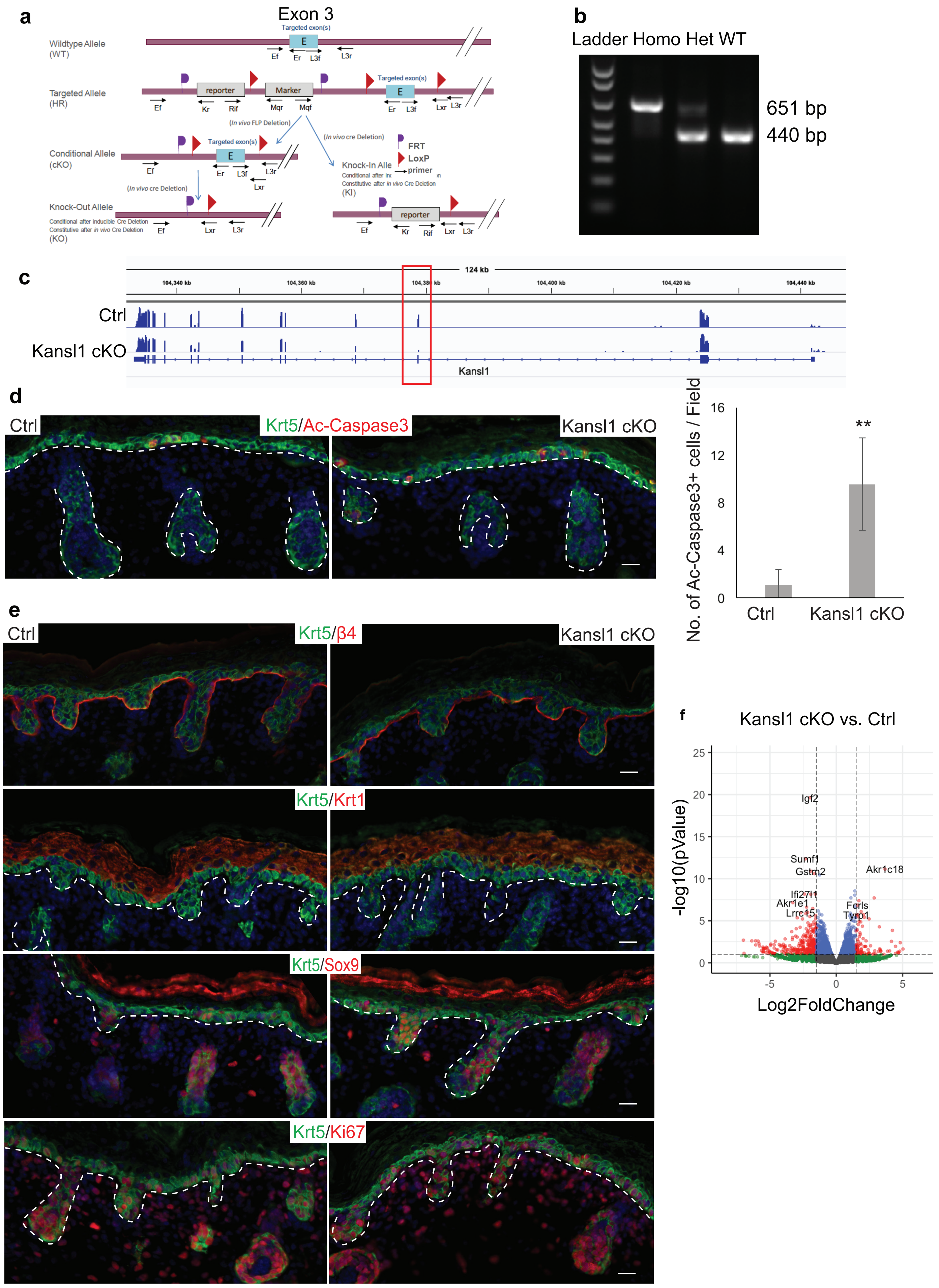

#### Extended Data Fig. 2

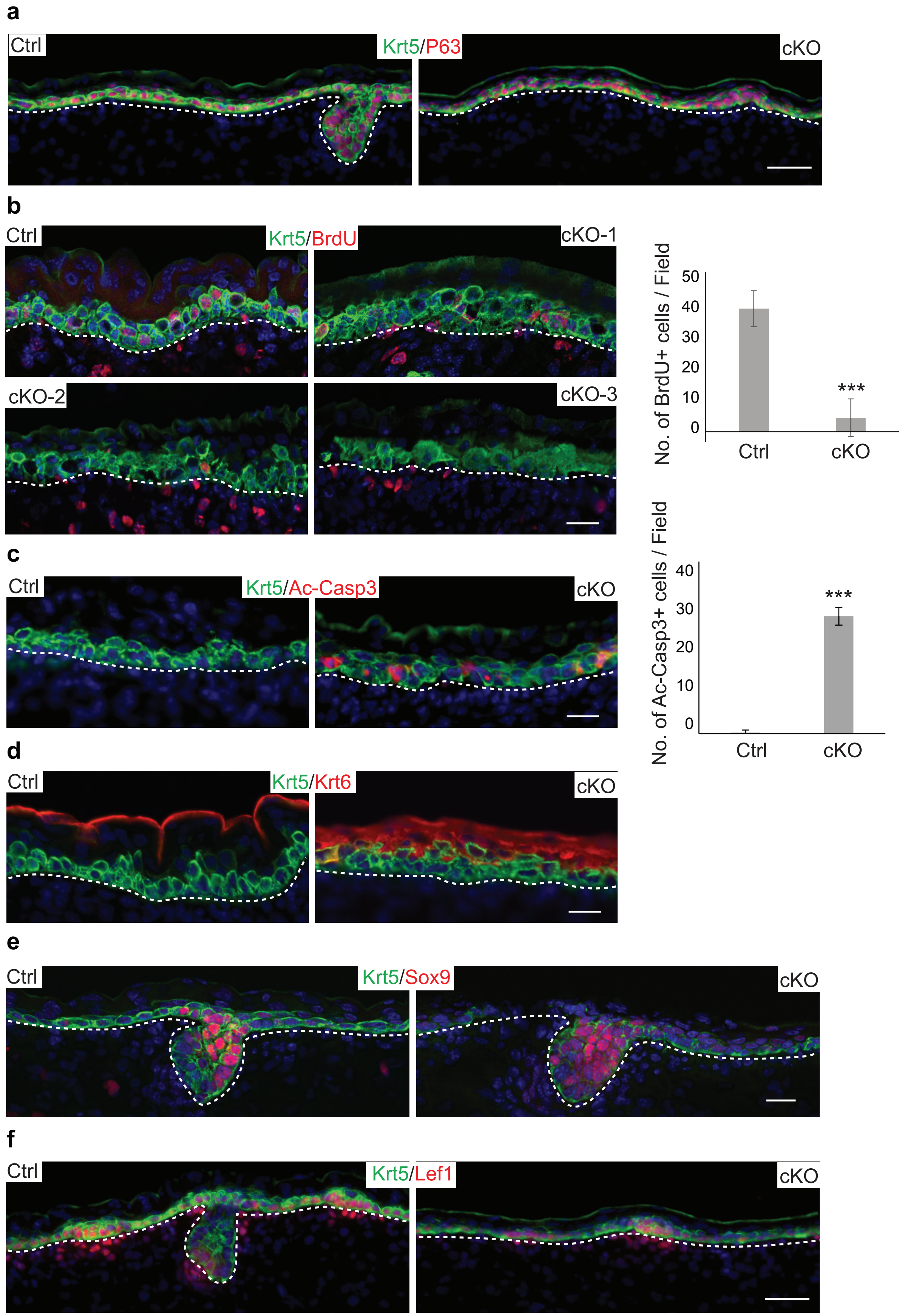

### Extended Data Fig. 3

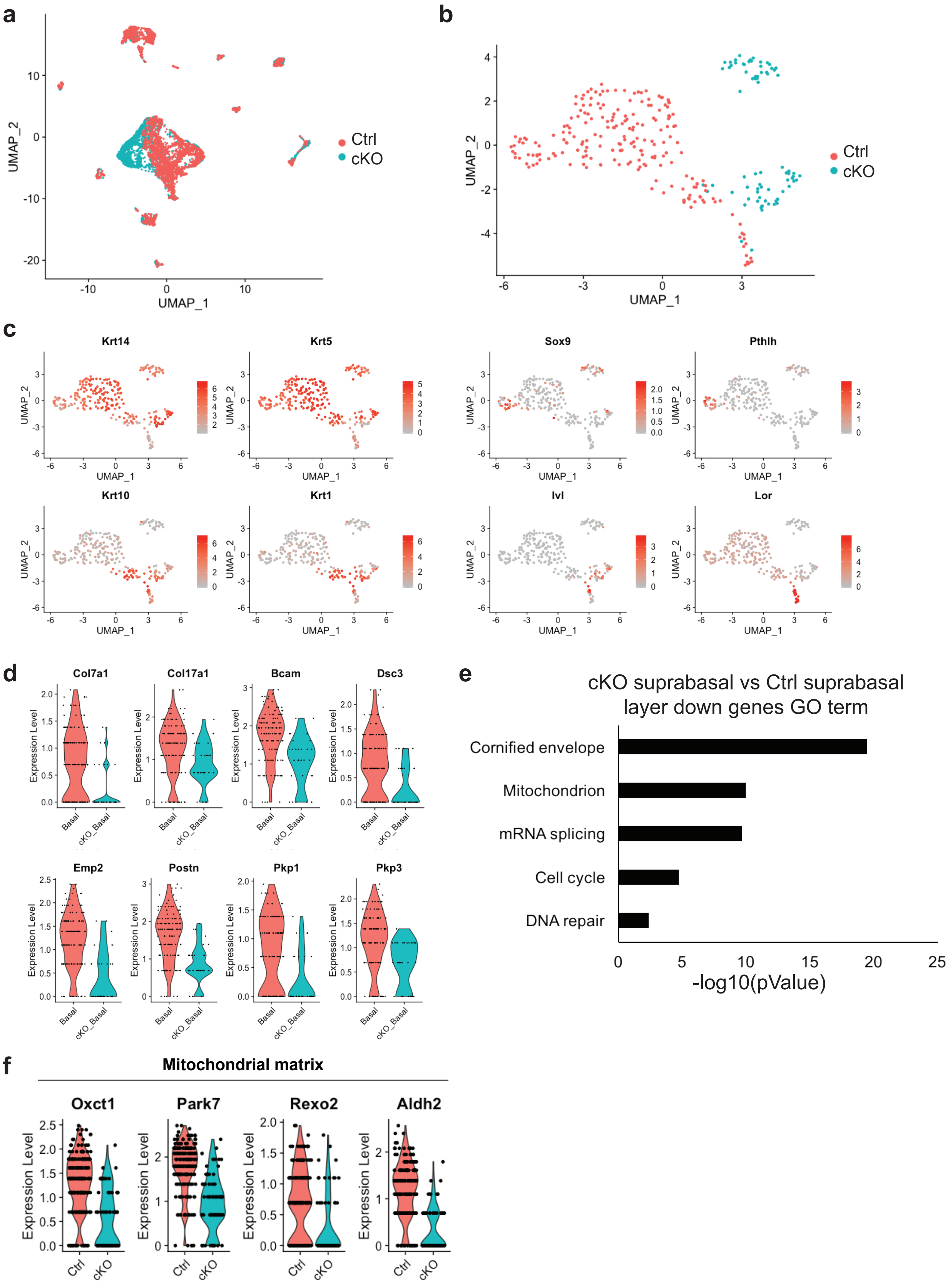

#### Extended Data Fig. 4

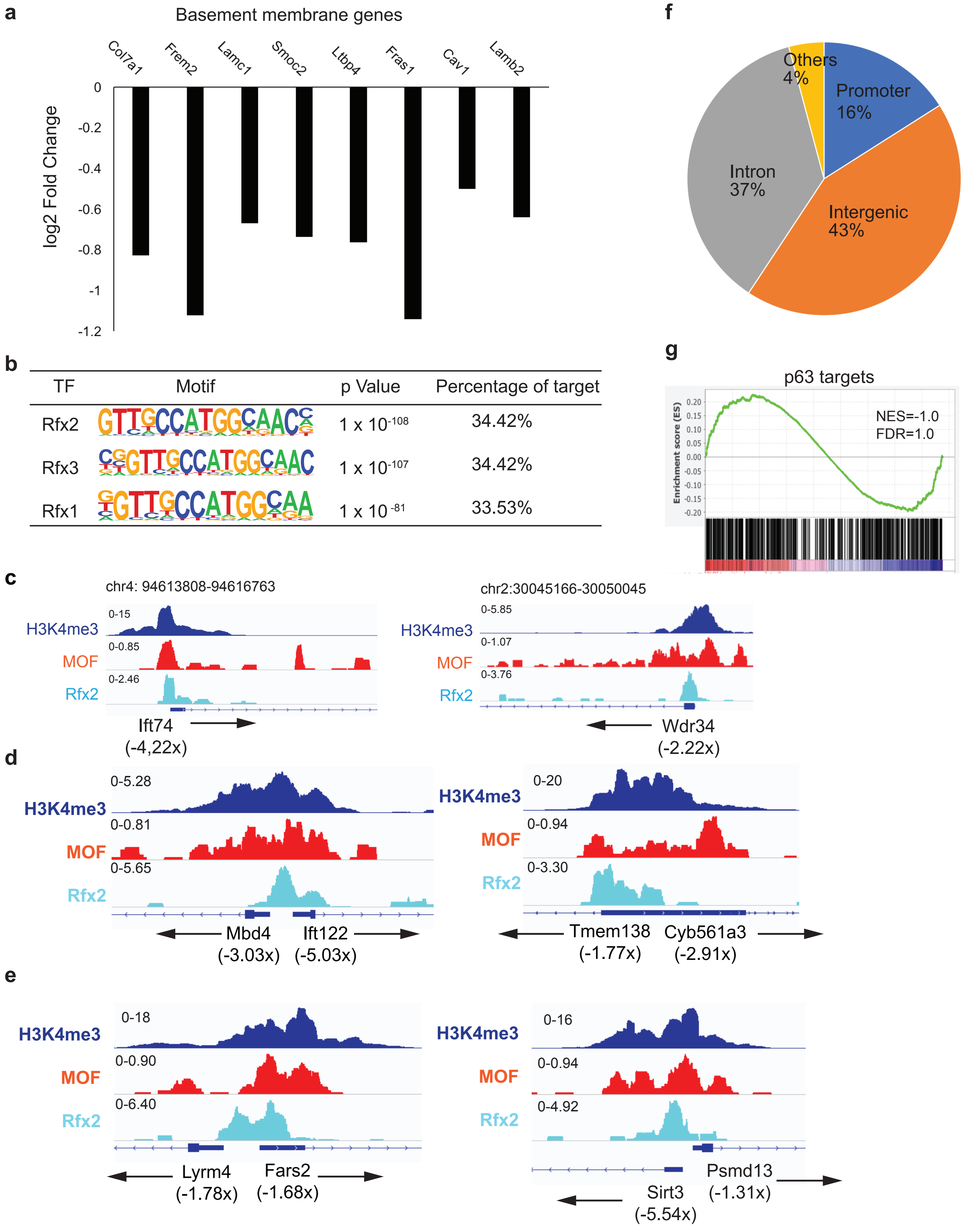

### Extended Data Fig. 5

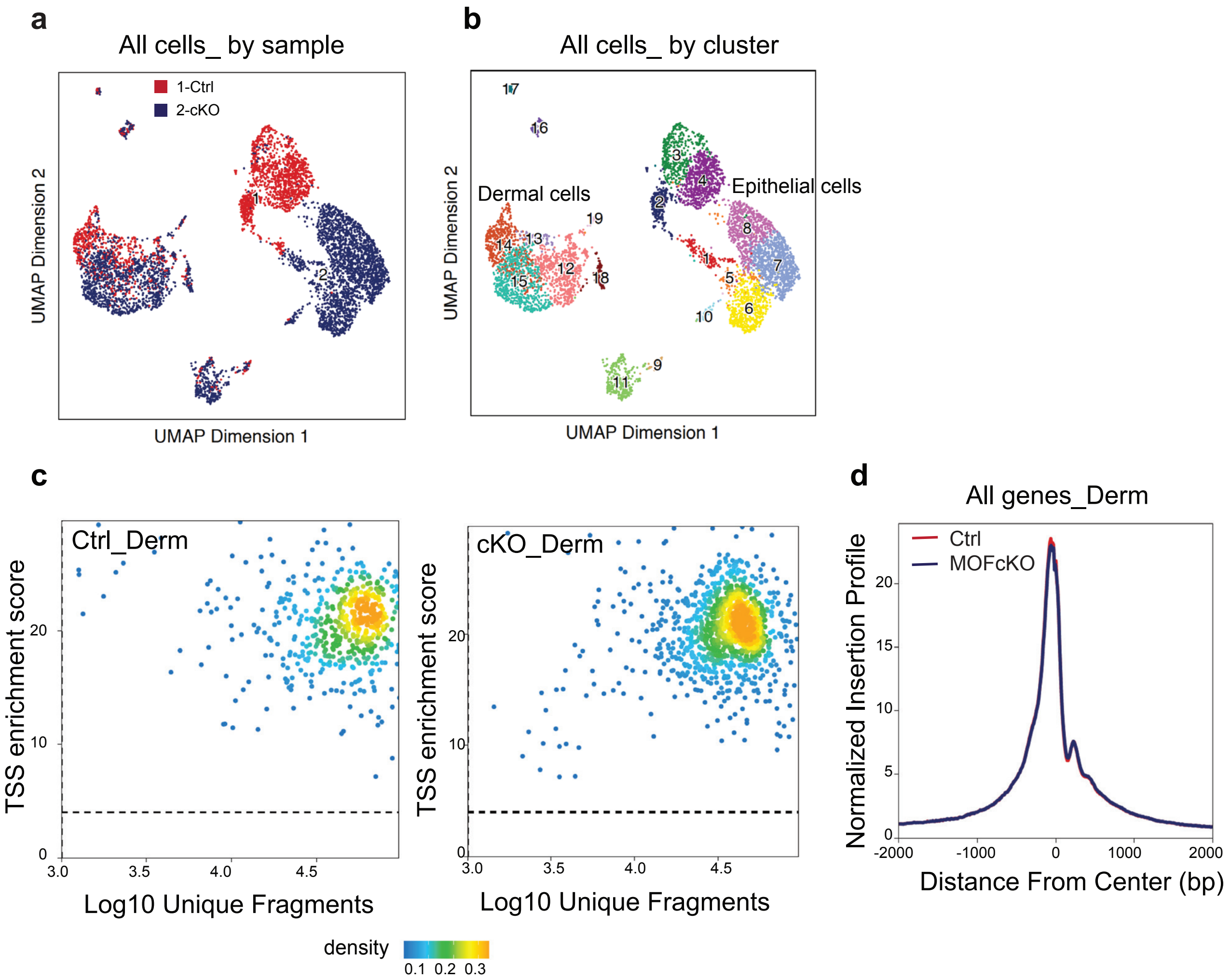

#### Extended Data Fig. 6

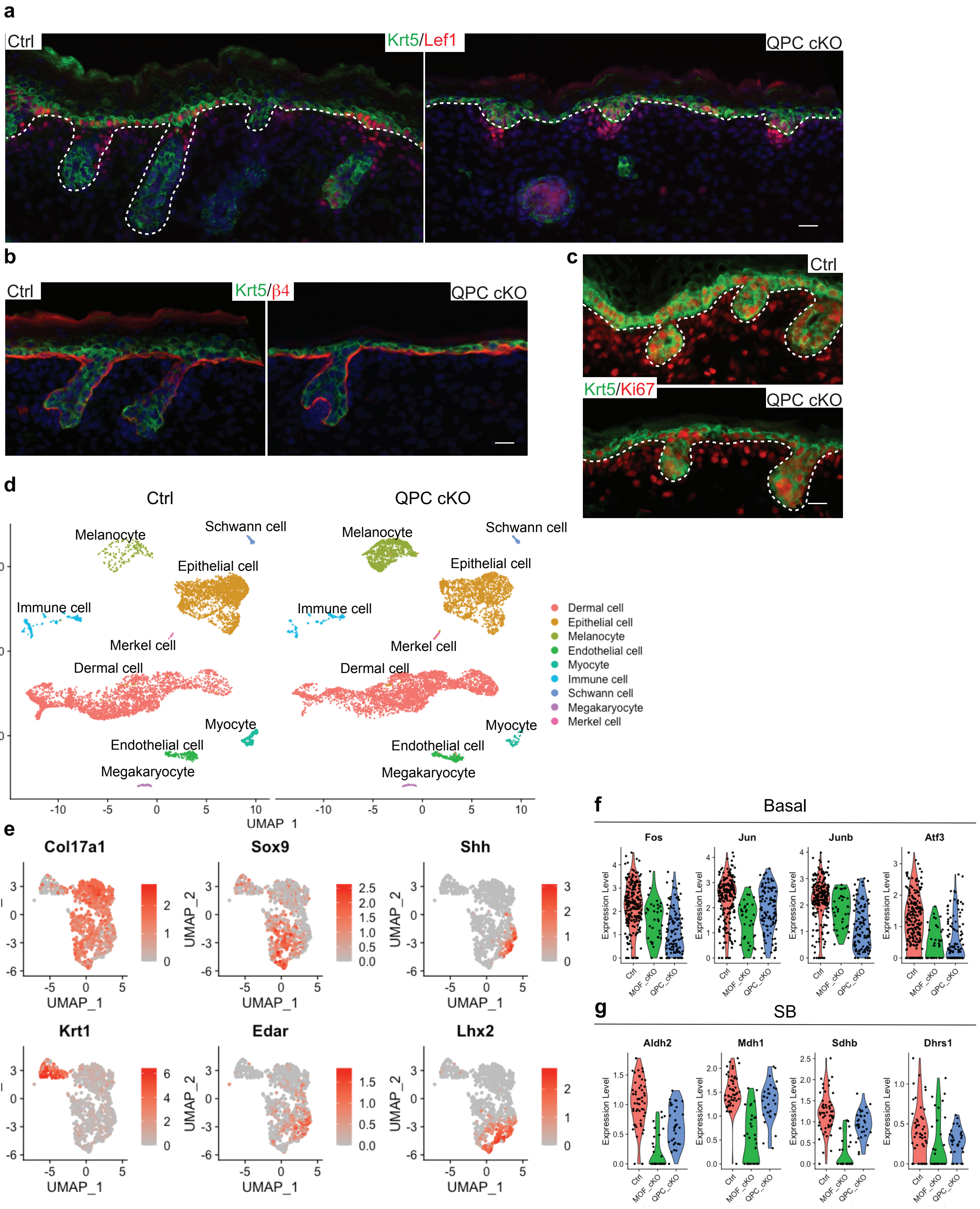
